# Targeting 3-mercaptopyruvate sulfurtransferase induces cancer stem cell death

**DOI:** 10.64898/2026.03.25.714276

**Authors:** Kelly Ascenção, Olivia Oravecz, Csaba Szabo

**Author notes:** Corresponding authors. &.

## Abstract

3-mercaptopyruvate sulfurtransferase (3-MST) is a mammalian enzyme that contributes to hydrogen sulfide and reactive sulfur species generation. Here we show that 3-MST is markedly upregulated in colorectal cancer stem cells (CSCs) and functions as a critical metabolic support mechanism for this therapy-resistant tumor cell population. CSCs exhibit low proliferation rate, high membrane rigidity and a metabolically restrained phenotype characterized by low oxidative phosphorylation rate, combined with a reduced rate of glycolysis. Genetic or pharmacological inhibition of 3-MST further suppresses cellular bioenergetics in CSCs, and this bioenergetic collapse impairs CSC proliferation, spheroid formation, migration and promotes cell death and attenuates tumor growth. Integrated transcriptomic, proteomic, metabolomic, and lipidomic analyses reveal extensive metabolic remodeling of the CSCs following 3-MST inhibition, including disruption of the glycolysis-TCA axis and marked remodeling of membrane lipid composition, including enrichment of ceramides and sphingolipids and increased incorporation of polyunsaturated phospholipids, resulting in increased membrane fluidity. 3-MST inhibition induced an activation of integrated stress pathways, proteotoxic stress responses and inflammatory signaling, linking the metabolic failure of CSCs to the induction of mixed-mode cell death. These findings identify 3-MST as a metabolic vulnerability in colorectal CSCs. Targeting this enzyme may be a translatable strategy to eliminate therapy-resistant tumor stem cell populations.

## MAIN

Hydrogen sulfide (H_2_S) is an endogenously produced gaseous signaling molecule that regulates multiple physiological and pathophysiological processes in mammalian cells [1–3]. In many cancers, H_2_S biosynthetic enzymes are upregulated, and their product, H_2_S supports tumor progression [4–7].

Cancer stem cells (CSCs) are a subpopulation of tumor cells with the capacity for self-renewal and differentiation and have been implicated in tumor propagation, therapeutic resistance and disease recurrence [8–11]. In colorectal cancer, accumulating evidence suggests that CSCs contribute to treatment failure and tumor relapse [12–14]. Targeting CSCs has therefore been proposed as a strategy to improve therapeutic outcomes; however, CSC populations exhibit pronounced resistance to conventional therapies, making their selective elimination a major challenge.

Recent proteomic analyses of human colon cancer cells have revealed distinct molecular features of the CSC population. Among the most prominent alterations is a marked upregulation of the H_2_S- and polysulfide-producing enzyme 3-mercaptopyruvate sulfurtransferase (3-MST) [15]. This enzyme, localized both in the mitochondria and the cytosol, is one of the major producers of reactive sulfur species (H_2_S, polysulfides) in mammals [2,3,16,17]. These findings raise the possibility that the 3-MST/H_2_S axis contributes to the maintenance and survival of colorectal CSCs. Here we investigated the functional role of 3-MST in colon CSCs. Our results identify 3-MST as a key metabolic support factor for these cells and reveal it as a previously unrecognized therapeutic vulnerability. Targeting 3-MST disrupts critical checkpoints in CSC metabolism to induce cell death, revealing a novel strategy for the selective elimination of therapy-resistant CSCs.

### 3-MST is upregulated in colon cancer CSCs

To determine whether sulfur metabolic pathways are altered in human tumors, we first analyzed paired colon tumor and adjacent control tissues. Multiple sulfur metabolism enzymes – including 3-MST, CARS2, SQR, CSE, TST, and ETHE1 – displayed increased expression in tumor samples relative to surrounding control tissue (Fig. 1A,B), with female predominance for the upregulation of CARS2, CBS and 3-MST (Extended Data Fig. 1). Similar patterns were observed across colon cancer cell lines, where several sulfur pathway enzymes were elevated compared with non-transformed colon epithelial cells (NCM356D) (Fig. 1C). As expected, cancer cells proliferated substantially faster than non-transformed epithelial cells (Fig. 1D,E) and the doubling time of these cell lines correlated strongly with the relative expression level of 3-MST (R² = 0.9736, p = 0.0133), suggesting a relationship between 3-MST abundance and proliferative capacity (Fig. 1F).

**Figure 1.**
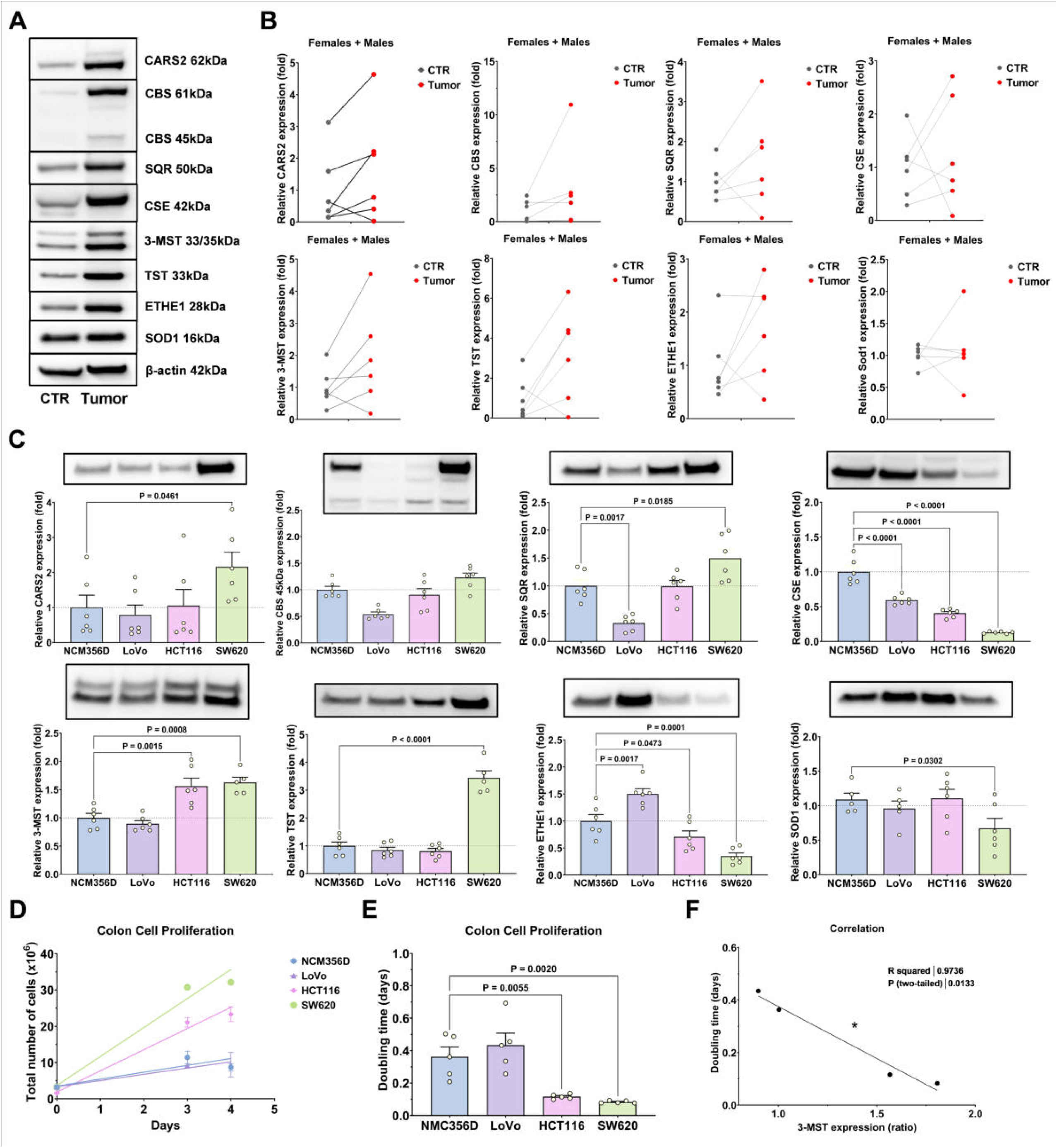
Upregulation of H_2_S–related enzymes in colorectal cancer and association with tumor cell proliferation. **(A)** Representative immunoblots showing protein expression of H_2_S-related enzymes and associated metabolic proteins in paired control (CTR) and colorectal tumor tissues. Proteins analyzed include CARS2, CBS (61 kDa and 45 kDa isoforms), SQR, CSE, 3-MST, TST, ETHE1 and SOD1; β-actin was used as a loading control. **(B)** Quantification of relative protein expression in paired control and tumor tissues. Each line represents paired samples from the same individual (N = 6; 3 females, 3 males). **(C)** Protein expression of the indicated enzymes in normal colon epithelial cells (NCM356D) and colorectal cancer cell lines (LoVo, HCT116 and SW620), determined by densitometric analysis of immunoblots normalized to β-actin. **(D)** Growth curves of NCM356D, LoVo, HCT116 and SW620 cells showing total cell number over time. **(E)** Quantification of cell doubling time for the indicated cell lines. **(F)** Correlation between relative 3-MST expression and cell doubling time across colon cell lines (R² = 0.9736, two-tailed P = 0.0133; N = 5). Data are presented as mean ± s.e.m from ≥5 independent experiments unless otherwise indicated.

### Cancer stem cells exhibit a metabolically quiescent state with altered sulfur metabolism and membrane properties

We next characterized CSCs derived from colon cancer lines and compared them with parental wild-type (WT) HCT116 colon cancer cells and with cells in a re-differentiated “rescue” state. CSC cultures formed large spheroid structures typical of stem-like growth, whereas WT cancer cells displayed a more differentiated morphology (Fig. 2A,B). Rescue cells retained a high clonogenic capacity despite re-exposure to differentiation conditions. Consistent with their stem-like phenotype [8–14], CSCs exhibited increased tumorigenic potential *in vivo*, as evidenced by larger tumor size compared with WT cancer cells (Fig. 2C). Flow-cytometric analysis confirmed enrichment of a subset of stem-associated surface markers in CSC populations (Extended Data Fig. 2A). As expected, the CSCs were markedly resistant to taxane-based chemotherapy while Rescue cells displayed an intermediate chemosensitivity phenotype (Fig. 2D; Extended Data Fig. 2B–D). Biophysical measurements revealed pronounced differences in membrane properties between the various cellular states investigated. In line with prior findings [18,19] CSC membranes were significantly more rigid than those of WT cancer cells, whereas Rescue cells showed membrane fluidity approaching WT levels but not fully restored (Fig. 2E). These changes were accompanied by altered expression of sulfur metabolism enzymes, including increased expression of 3-MST, CSE, SQR, TST, and ETHE1 in CSCs (Fig. 2F). Notably, Rescue cells maintained high 3-MST expression levels, suggesting that sustained 3-MST expression is a feature of the CSC phenotype. Consistent with their metabolic remodeling, CSCs produced higher levels of H_2_S compared with WT, whereas Rescue cells showed a partial retention of increased H_2_S production (Extended Data Fig. 2E,F).

**Figure 2.**
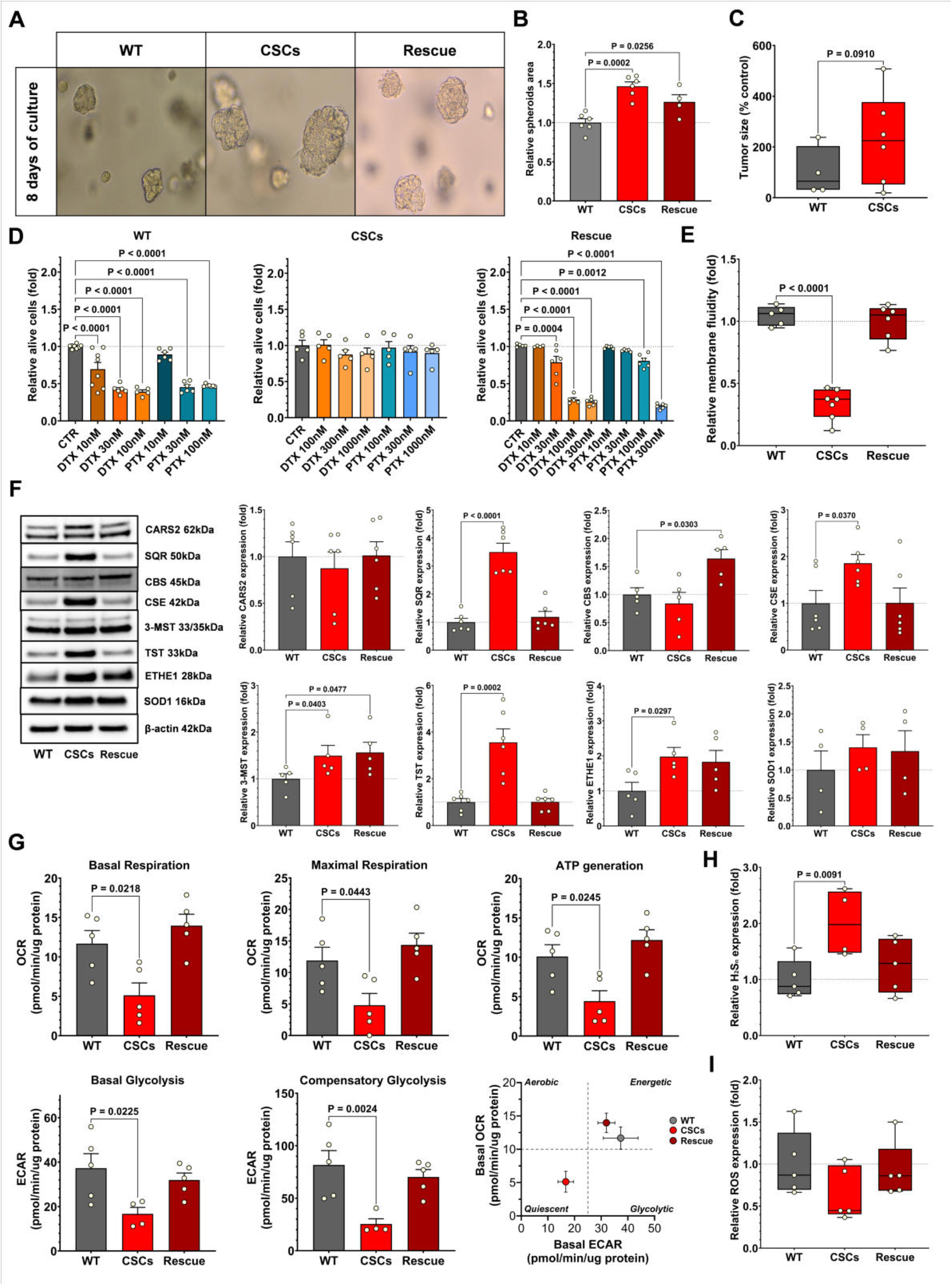
Cancer stem cell–associated metabolic reprogramming and H_2_S pathway alterations. **(A)** Representative images of spheroids formed after 8 days of culture from HCT116 wild-type cells (WT), HCT116-derived cancer stem cells (CSCs), and re-differentiated CSCs (Rescue). **(B)** Quantification of relative spheroid area. **(C)** Tumor growth expressed as percentage of control in WT- and CSC-derived tumors (WT, n = 4 males; CSCs, n = 6 males). **(D)** Cell viability following treatment with the microtubule-targeting chemotherapeutic agents docetaxel (DTX) and paclitaxel (PTX). Viable cells were normalized to untreated controls. **(E)** Relative membrane fluidity in WT, CSCs and Rescue cells. **(F)** Representative immunoblots and densitometric quantification of H_2_S-related enzymes and associated metabolic proteins in WT, CSCs and Rescue cells, including CARS2, SQR, CBS, CSE, 3-MST, TST, ETHE1 and SOD1; β-actin was used as a loading control. **(G)** Bioenergetic profiling of WT, CSCs and Rescue cells measured by extracellular flux analysis, including basal respiration, maximal respiration, ATP production, basal glycolysis and compensatory glycolysis. **(H)** Relative levels of reactive sulfur species (H_2_S_n_) in WT, CSCs and Rescue cells. **(I)** Relative reactive oxygen species (ROS) levels in WT, CSCs and Rescue cells. Data are presented as mean ± s.e.m from ≥4 independent experiments unless otherwise indicated.

To determine whether these phenotypic differences were associated with metabolic changes, we performed extracellular flux analysis. CSCs exhibited a markedly suppressed metabolic state, characterized by reduced mitochondrial respiration and glycolytic activity compared with WT cancer cells, whereas Rescue cells displayed metabolic activity closer to WT levels (Fig. 2G). Consistent with their quiescent phenotype, bioenergetic parameters were lower in CSCs than in WT HCT116 cells: basal respiration, maximal respiration, ATP production, basal glycolysis, and compensatory glycolysis were low in CSCs (Fig. 2G). In parallel, CSCs displayed increased levels of reactive sulfur species (H_2_S_n_) relative to WT cells, whereas Rescue cells exhibited intermediate H_2_S_n_ levels (Fig. 2H). Conversely, ROS levels tended to be lower in CSCs compared with WT cells, with Rescue cells showing intermediate levels (Fig. 2I). CSCs showed a trend toward increased telomere length relative to WT cells, consistent with their stem-like properties (Extended Data Fig. 2G).

### Reduced 3-MST expression impairs proliferation, stemness, and migration

To assess the functional role of 3-MST, we generated MPST-deficient colon cancer cells (Fig. 3A–C). MPST-deficient cells displayed subtle morphological changes compared with WT cells (Fig. 3A). Reduced 3-MST expression significantly decreased the proliferative capacity of WT colon cancer cells (Fig. 3B,C). MPST deficiency also affected the expression of other sulfur metabolism enzymes. In WT cells, reduced 3-MST expression was associated with decreased levels of CBS and TST, whereas other components of the sulfur metabolic network were only modestly affected (Extended Data Fig. 3A,B). In contrast, CSCs exhibited increased ETHE1 levels together with modest alterations in other sulfur metabolism enzymes, suggesting selective adaptation within the sulfur metabolic network (Extended Data Fig. 3C,D). In CSCs, reduced 3-MST expression exerted a more pronounced effect. MPST-deficient CSC cultures showed impaired spheroid growth and markedly reduced expansion over time compared with control CSCs (Fig. 3D-G), accompanied by a significant increase in doubling time (Fig. 3G). Furthermore, spheroid formation (a key indicator of cancer stemness) was significantly reduced in MPST-deficient CSCs (Fig. 3H,I). Migration capacity was also impaired (Fig. 3J–L).

**Figure 3.**
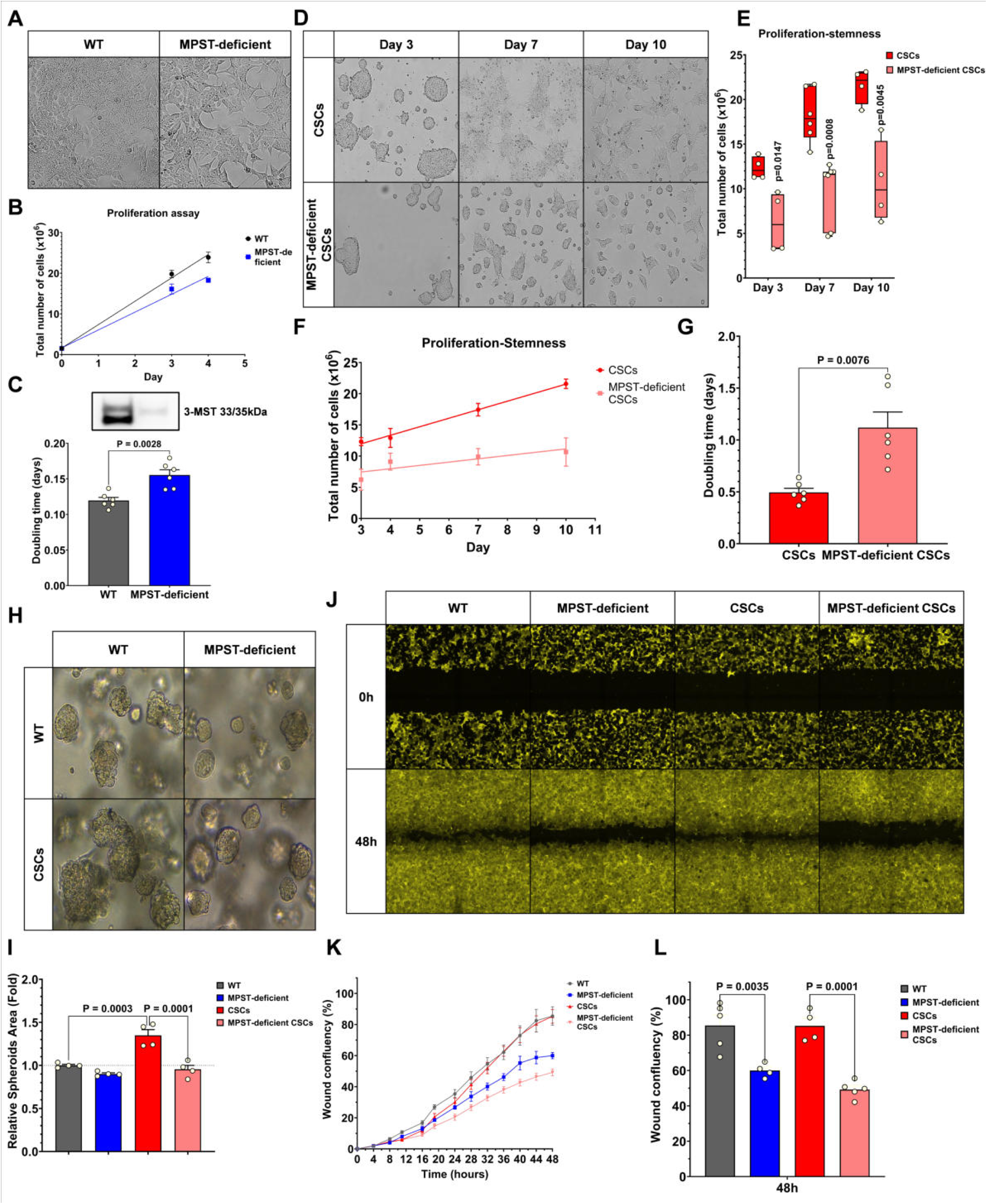
Reduced 3-MST expression impairs proliferation, stemness and migratory capacity of colorectal cancer cells. **(A)** Representative phase-contrast images of HCT116 wild-type (WT) and MPST-deficient cells showing morphological changes associated with reduced 3-MST expression. **(B)** Proliferation assay showing total cell number over time in WT and MPST-deficient cells. Data are presented as mean ± s.e.m. from 3 independent experiments. **(C)** Quantification of cell doubling time in WT and MPST-deficient cells. Inset shows a representative immunoblot confirming reduced 3-MST protein expression. **(D)** Representative images of cancer stem cell cultures (CSCs) and MPST-deficient CSCs during expansion at different time points (days 3, 7 and 10). **(E)** Quantification of CSC expansion showing total cell number at days 3, 7 and 10. **(F)** Growth curves of CSCs and MPST-deficient CSCs. **(G)** Quantification of doubling time in CSCs and MPST-deficient CSCs. **(H)** Representative images of spheroids formed by WT cells, MPST-deficient cells, CSCs and MPST-deficient CSCs. **(I)** Quantification of relative spheroid area. **(J)** Representative images from wound-healing assays showing migration at 0 h and 48 h in WT, MPST-deficient WT, CSCs and MPST-deficient CSCs. **(K)** Time-course quantification of wound closure. **(L)** Quantification of wound confluency at 48 h. Data are presented as mean ± s.e.m. from ≥4 independent experiments unless otherwise indicated.

### 3-MST inhibition preferentially targets cancer stem cells: disruption of bioenergetics, suppression of clonogenicity, increases in membrane fluidity and induction of cell death

To determine whether pharmacological inhibition of 3-MST recapitulates the effects of reduced 3-MST expression, we treated cells with the 3-MST inhibitor HMPSNE [16,17]. HMPSNE treatment disrupted spheroid growth in both WT and CSC cultures, with a more pronounced effect in CSCs (Fig. 4A,B). HMPSNE treatment significantly reduced viability in both WT and CSCs, with CSCs exhibiting the greatest sensitivity (Fig. 4C, Extended Data Fig. 4A). Inhibition of 3-MST also suppressed cell viability and induced death of SW620 CSCs and LoVo CSCs (Extended Data Fig. 4B,C).

**Figure 4.**
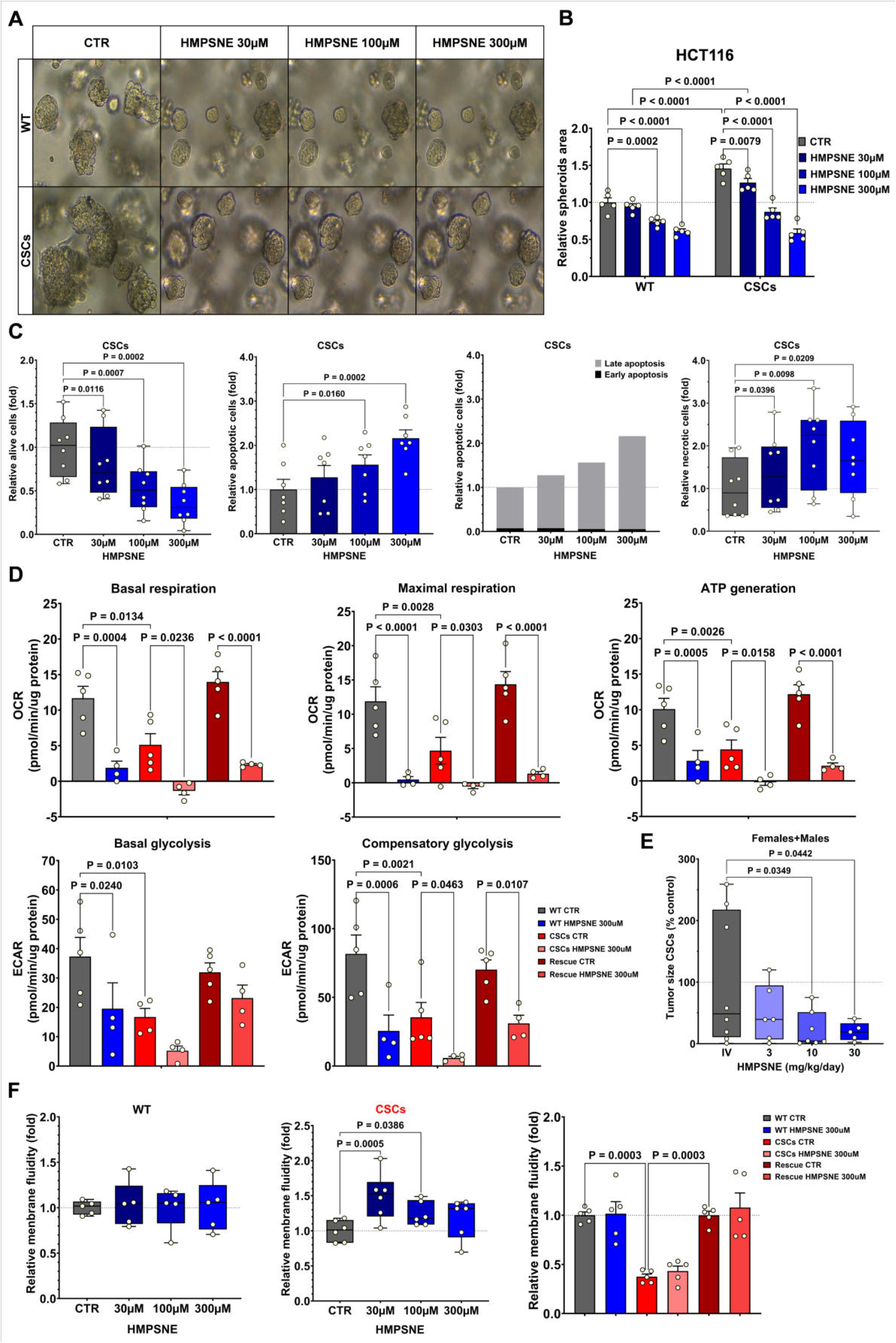
Pharmacological inhibition of 3-MST suppresses cancer stem cell viability, metabolism and tumor growth. **(A)** Representative images of spheroids formed by HCT116 wild-type (WT) cells and cancer stem cells (CSCs) treated with increasing concentrations of the 3-MST inhibitor HMPSNE (30-300 μM) compared with untreated controls (CTR). **(B)** Quantification of relative spheroid area in WT and CSCs cultures following HMPSNE treatment. **(C)** Quantification of cell viability and cell death parameters (apoptosis and necrosis) in CSCs treated with HMPSNE. Early and late apoptotic populations are shown separately. **(D)** Bioenergetic profiling of WT, CSCs and Rescue cells treated with HMPSNE (300 μM) measured by extracellular flux analysis, including basal respiration, maximal respiration, ATP production, basal glycolysis and compensatory glycolysis. **(E)** *In vivo* tumor growth in an HCT116 CSCs-derived xenograft model following administration of HMPSNE (3-30 mg kg⁻¹ day⁻¹). **(F)** Membrane fluidity measurements in WT, CSCs and Rescue cells following HMPSNE treatment. Data are presented as mean ± s.e.m from ≥4 independent experiments unless otherwise indicated. For *in vivo* experiments (E): CTR, n = 8; 3 mg kg⁻¹, n = 6; 10 mg kg⁻¹, n = 7; 30 mg kg⁻¹, n = 5.

HMPSNE treatment also altered the expression of enzymes involved in sulfur metabolism. Inhibition of 3-MST reduced the expression of several sulfur-metabolic enzymes including 3-MST, SQR, TST, and ETHE1, indicating broader disruption of the sulfur metabolic network (Extended Data Fig. 4D). As expected, the 3-MST inhibitor produced a concentration-dependent inhibition of H_2_S generation in WT, CSC as well as Rescue cells (Extended Data Fig. 4E,F).

Extracellular flux analysis revealed that HMPSNE treatment profoundly suppresses cellular bioenergetics. 3-MST inhibition reduced both oxidative phosphorylation and glycolysis in WT cancer cells, but a detectable residual activity remained. In contrast, in CSCs (which exhibited lower baseline oxidative phosphorylation and glycolysis than WT cancer cells) 3-MST inhibition further suppressed all of the measured bioenergetic parameters, resulting in a near-complete suppression of glycolysis as well as oxidative phosphorylation. Rescue cells displayed a response similar to WT cells, with residual bioenergetic activity preserved (Fig. 4D). These effects of 3-MST inhibition are consistent with the supporting role of this enzyme to maintain glycolysis and oxidative phosphorylation in various cell types [4], but are functionally more critical in CSCs, which have a low baseline cellular bioenergetic activity.

The metabolic collapse induced by HMPSNE was associated with antitumor effects *in vivo*. In a murine xenograft model derived from colon CSCs, systemic HMPSNE treatment markedly reduced tumor growth (Fig. 4E).

Inhibition of 3-MST reduced cellular ROS levels in WT and Rescue cells, but not CSCs (Extended Data Fig. 5A). In CSCs, HMPSNE treatment altered the cell-surface marker profile, with increases in CD24 and LGR5, decreased CD133 expression, and reduced expression across several other markers (Extended Data Fig. 5B). In addition 3-MST inhibition significantly altered the biophysical properties of cellular membranes. In CSCs, which display highly rigid membranes, 3-MST inhibition increased membrane fluidity (Fig. 4F). Furthermore, HMPSNE treatment was associated with an increase in telomere length in CSCs (Extended Data Fig. 5C).

### 3-MST inhibition induces transcriptional, proteomic, metabolomic and lipidomic remodeling in CSCs

Transcriptomic, proteomic, metabolomic, and lipidomic analyses (Supplementary Data 1-3) were conducted to define the biochemical consequences of 3-MST inhibition in CSCs. Principal component analysis (PCA) revealed clear separation between WT cancer cells, CSCs, and HMPSNE-treated CSCs, indicating substantial global differences in gene expression, protein abundance, and metabolic state among these conditions (Fig. 5A). Rescue cells largely overlapped with WT cells across transcriptomic, metabolic and lipidomic profiles, whereas the proteome remained distinctly altered (Supplementary Data 5). Volcano plot analysis, heatmaps, Revigo and LION enrichment analysis further demonstrated widespread remodeling of the proteome, metabolome and lipidome of CSCs compared with WT cancer cells, as well as after pharmacological inhibition of 3-MST (Fig. 5B,C; Fig. 6. A,B; Supplementary Data 1-3). Many altered proteins were linked to central carbon metabolism and lipid biosynthesis, consistent with metabolomic and lipidomic changes (Fig. 5D; Fig. 6C; Supplementary Data 1,2; Table 1).

**Figure 5.**
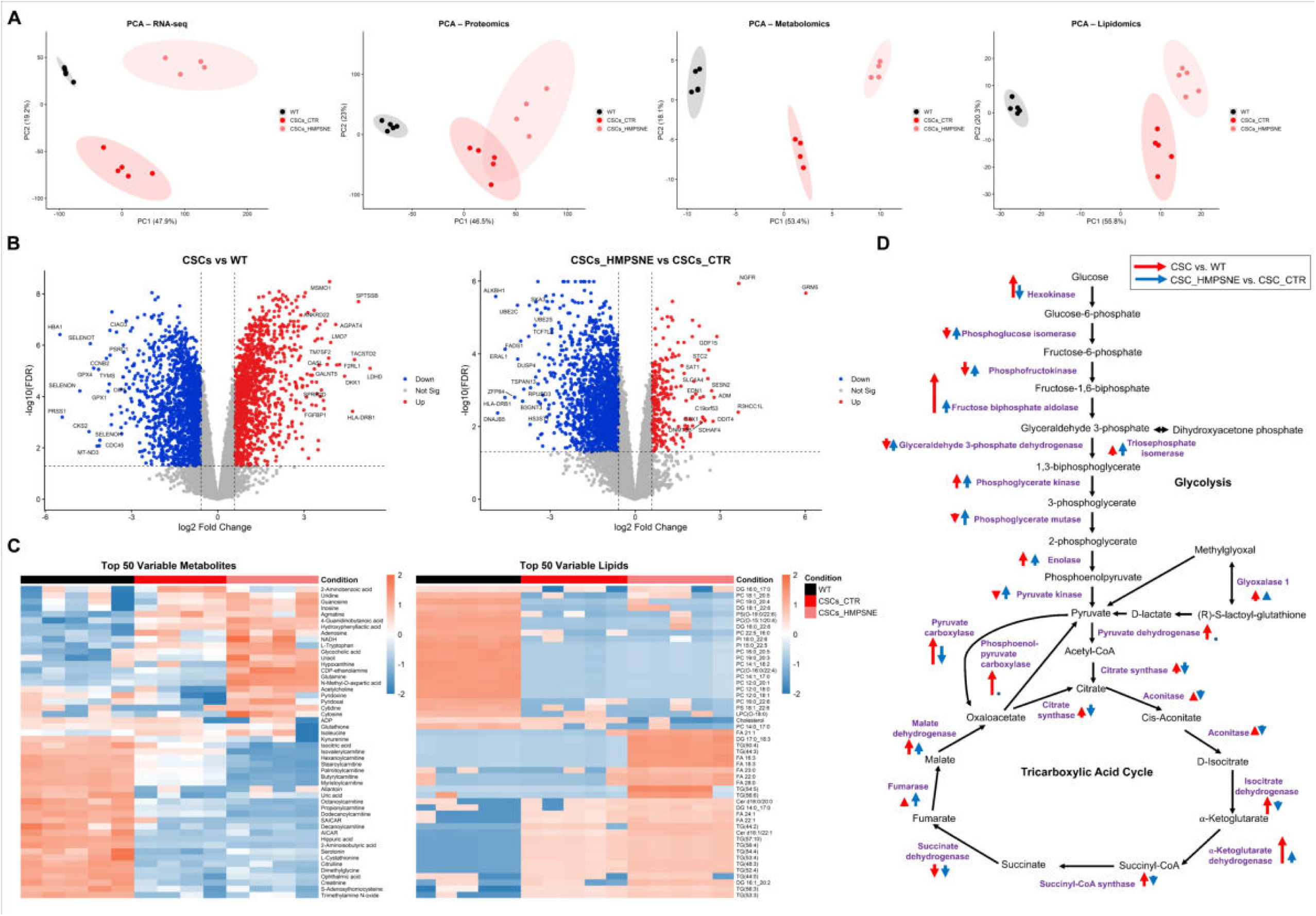
Integrative multi-omics profiling reveals proteomic, metabolic and lipidomic reprogramming in cancer stem cells and its modulation by 3-MST inhibition. **(A)** Principal Component Analysis (PCA) of RNA-seq, proteomics, metabolomics and lipidomics datasets analyzed independently, showing separation between wild-type (WT), cancer stem cell controls (CSCs_CTR) and HMPSNE-treated CSCs (CSCs_HMPSNE). Each dot represents an independent biological replicate; ellipses indicate clustering of samples within each group. **(B)** Volcano plots of differential protein expression. Left: CSCs compared with WT cells. Right: CSCs treated with HMPSNE compared with untreated CSC controls. Red and blue dots indicate significantly upregulated and downregulated proteins, respectively (|fold change| ≥1.5; FDR ≤0.05), whereas grey dots represent non-significant changes. The top 15 upregulated and top 15 downregulated proteins are labeled. **(C)** Heat maps of the 50 most variable metabolites (left) and lipid species (right) across WT, CSCs_CTR and CSCs_HMPSNE samples. Rows represent metabolites or lipid species and columns represent individual samples. Colors indicate relative abundance (z-score normalized values). **(D)** Schematics of lipid and sterol biosynthesis pathways. Red arrows indicate enzymes differentially expressed between CSCs and WT cells, whereas blue arrows denote enzymes altered following HMPSNE treatment.

**Figure 6.**
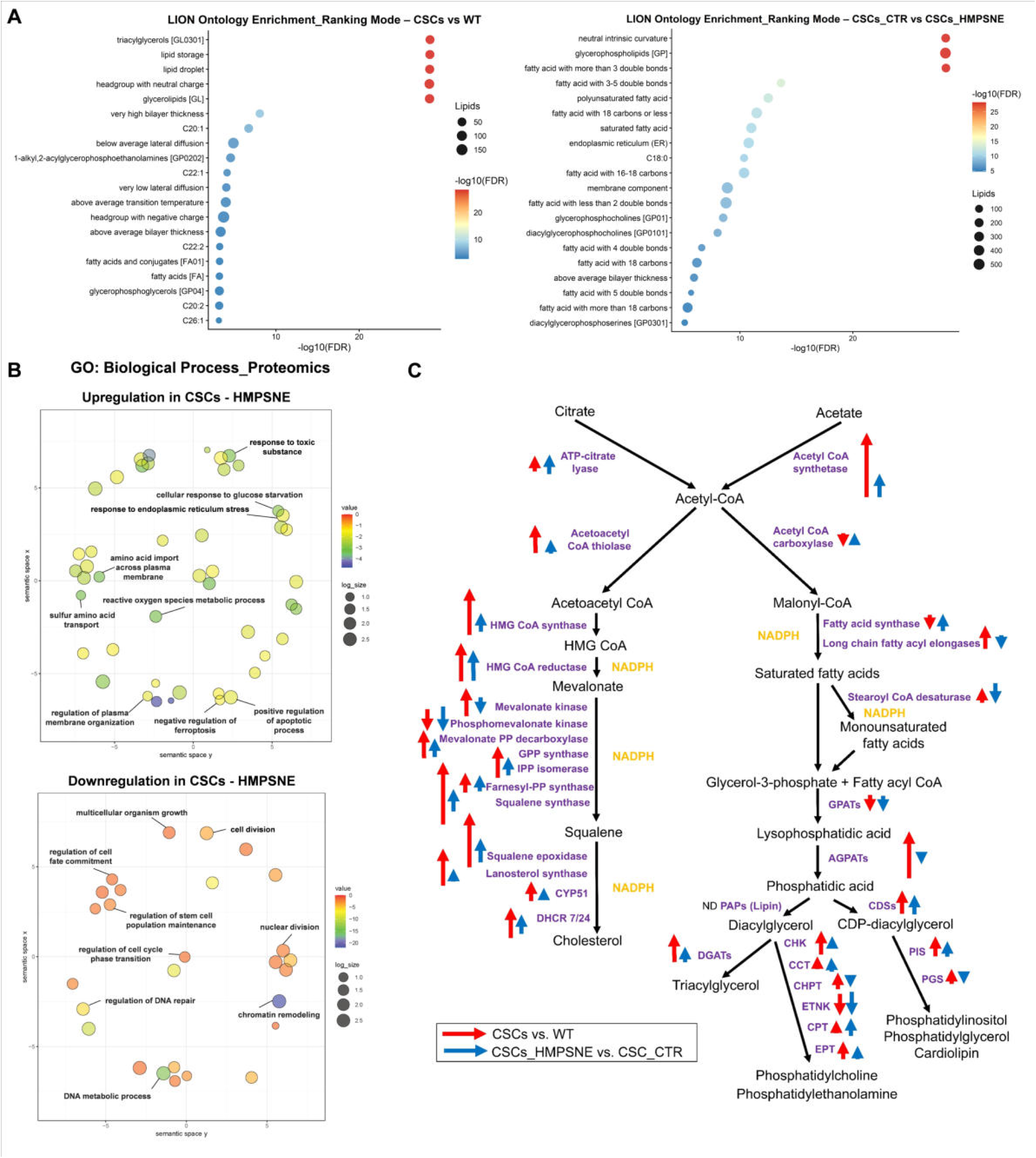
Integrative multi-omics profiling reveals proteomic, metabolic and lipidomic reprogramming in cancer stem cells and its modulation by 3-MST inhibition. **(A)** LION (Lipid Ontology) enrichment analysis identifying lipid classes enriched in CSCs relative to WT (left) and in CSC controls relative to HMPSNE-treated CSCs (right). Dot size represents the number of lipid species contributing to enrichment and color indicates enrichment significance (−log10 FDR). **(B)** Gene ontology (GO) biological process enrichment derived from proteomics data in HMPSNE-treated CSCs. Semantic similarity plots summarize significantly upregulated (top) and downregulated (bottom) biological processes relative to CSC controls. Point size corresponds to the number of associated proteins and color indicates enrichment significance (−log10 FDR). **(C)** Schematics of glycolysis and the tricarboxylic acid (TCA) cycle. Red arrows indicate enzymes differentially expressed between CSCs and WT cells, whereas blue arrows denote enzymes altered following HMPSNE treatment.

Comparison of CSCs with WT cells revealed coordinated changes in enzymes involved in glycolysis and the TCA cycle. Several enzymes regulating glucose utilization and mitochondrial carbon entry were increased in CSCs, including hexokinase, pyruvate carboxylase, PCK2, pyruvate dehydrogenase, isocitrate dehydrogenase, α-ketoglutarate dehydrogenase, and malate dehydrogenase, whereas succinate dehydrogenase, which links the TCA cycle to the electron transport chain, was reduced (Fig. 5D). Despite this expanded enzymatic capacity, functional measurements demonstrated that CSCs operate at reduced metabolic throughput compared with WT tumor cells, as measured by extracellular flux analysis (see above). Metabolomic analysis supported these findings and revealed depletion of key intermediates in central carbon metabolism, including reductions in pyruvate and upper TCA intermediates such as citrate, cis-aconitate, and isocitrate (Fig. 5D).

Inhibition of 3-MST significantly perturbed the metabolic architecture of CSCs. Proteomic analysis showed decreases in several enzymes involved in mitochondrial carbon entry and early TCA-cycle reactions, including pyruvate carboxylase, citrate synthase, aconitase, and isocitrate dehydrogenase (Fig. 5D). Consistent with these changes, metabolomic analysis revealed further depletion of upper TCA intermediates. Together with a simultaneous suppression of oxidative phosphorylation and glycolysis after 3-MST inhibition in CSCs, pyruvate and lactate levels increased following 3-MST inhibition (Supplementary Data 2; Table 1), suggesting diversion of carbon away from mitochondrial oxidation due to impaired entry into the TCA cycle (Fig. 5D; Table 1).

In addition to the pronounced suppression of mitochondrial function, the proteomic data also revealed alterations in proteins associated with mitochondrial structure and protein homeostasis after 3-MST inhibition (Table 1). Several proteins involved in mitochondrial ribosome assembly and mitochondrial RNA processing were markedly reduced following HMPSNE treatment of the CSCs, including the mitochondrial rRNA chaperone ERAL1 and members of the pseudouridine synthase family RPUSD. In addition to these factors, multiple mitochondrial ribosomal proteins were also decreased, indicating a broader suppression of the mitochondrial translation machinery. Components of respiratory complex I were similarly affected, with reduced abundance of numerous subunits, and several members of the mitochondrial redox protein family FDX. These changes suggest that 3-MST inhibition disrupts mitochondrial RNA maturation, mitochondrial protein synthesis, and respiratory chain integrity, contributing to the impairment of mitochondrial maintenance in CSCs.

Proteomic analysis revealed reduced abundance of multiple regulators of cell-cycle progression and mitosis, including CKS1B, UBE2C, UBE2S, SKA3, SPC25, CCNB2, CKS2, PCLAF and PTTG1 after 3-MST inhibition, consistent with suppression of proliferative programs and a shift toward growth arrest.

3-MST inhibition also induced pronounced remodeling of cellular lipid composition. Lipidomic analysis revealed widespread remodeling across multiple lipid classes, including glycerophospholipids and sphingolipids (Fig. 5C; Supplementary Data 2). Additional lipidomic changes were observed in lipid classes associated with membrane organization and curvature, including lysoglycerophospholipids, monoacylglycerophospholipids, diacylglycerols, and phospholipids containing very-long-chain or polyunsaturated fatty acids (Supplementary Data 2). These alterations were accompanied by changes in lipid metabolic pathways, including sterol and phospholipid metabolism, which likely influenced membrane physical properties.

The proteomic profile of CSCs indicated coordinated alterations in enzymes involved in sterol and fatty-acid metabolism (Fig. 5B, Table 1). Multiple enzymes in the mevalonate and sterol synthesis pathways, including HMGCR, HMGCS1, FDFT1, SQLE, DHCR7, DHCR24, LSS, and MSMO1, were increased in CSCs. At the same time, the polyunsaturated fatty acid-producing enzyme FADS1 was reduced, while several fatty-acid elongases were increased. Additional changes in phospholipid remodeling enzymes, including members of the AGPAT family, LPCAT enzymes, and DGAT1 (Table 1), were also observed. These coordinated proteomic changes indicate activation of anabolic lipid metabolism in CSCs, providing a mechanistic basis for increased membrane synthesis and structural remodeling observed in lipidomic analyses. Treatment of CSCs with the 3-MST inhibitor induced several changes in these pathways. Early steps of the mevalonate pathway were partially reduced, including decreases in PMVK. In parallel, enzymes involved in phospholipid remodeling and fatty-acid desaturation were reduced, including marked suppression of FADS1 and SCD, consistent with impaired generation of polyunsaturated and monounsaturated fatty acids. HMPSNE treatment decreased major membrane phospholipids and cholesterol while increasing free fatty acids and diacylglycerols, indicating disruption of membrane homeostasis (Supplementary Data 2; Table 1). These alterations were accompanied by reduced membrane unsaturation (Supplementary Data 2) and increased membrane rigidity (see above), reflecting substantial remodeling of membrane lipid composition following 3-MST inhibition.

Proteins that bind membrane phospholipids and regulate lipid-dependent signaling play key roles in membrane organization and lipid oxidation processes. Among these, the phosphatidylethanolamine-binding protein PEBP1 functions as a lipid-binding scaffold that can facilitate lipoxygenase-mediated oxidation of membrane phospholipids. In the proteomic analysis, PEBP1 levels were modestly reduced in CSCs relative to WT cells but increased following treatment of CSCs with the 3-MST inhibitor (Table 1), suggesting enhanced phospholipid oxidation capacity. In parallel, the expression of stearoyl-CoA desaturase 1 (SCD), which generates monounsaturated fatty acids that protect membranes from lipid peroxidation – a known enzyme involved in maintaining stem cell rigidity [18,19] – was reduced after HMPSNE treatment. Increased PEBP1 and decreased SCD would be expected to favor lipid-peroxide formation, thereby promoting membrane disorder and increased susceptibility to ferroptotic lipid damage in CSCs [20]. Enrichment analyses further identified ferroptosis-related signatures (Supplementary Data 2). The increased pro-oxidant lipid species, reduced protective lipid desaturation (SCD↓), enhanced phospholipid oxidation capacity (PEBP1↑), and pathway-level enrichment indicate that 3-MST inhibition promotes a ferroptosis-prone membrane state in CSCs.

In addition to these metabolic and lipid alterations, proteomic analysis revealed coordinated changes in proteins involved in cystine transport, redox homeostasis and amino-acid stress responses, including SLC7A11, SLC3A2, PHGDH, PSAT1, TXNRD1, SRXN1 and SESN2, which were increased following HMPSNE treatment (Table 1). These changes indicate activation of compensatory antioxidant defense mechanisms. In parallel, GPX4 was reduced, whereas SAT1 and HMOX1 were strongly induced, consistent with increased oxidative and lipid-peroxidation stress and impaired detoxification of lipid peroxides. Thus, 3-MST inhibition promotes lipid oxidative damage and ferroptotic vulnerability. Enrichment analyses further identified pathways related to the integrated stress response, endoplasmic reticulum (ER) stress and misfolded-protein responses, indicating activation of ER stress and the unfolded protein response in CSCs (Supplementary Data 2). Proteomic data further supported activation of proteostasis pathways, with increased levels of stress-associated proteins including SQSTM1 and DNAJB9. Consistent with this, Gene Ontology enrichment analysis revealed activation of stress-related biological processes, including responses to toxic substances, ER stress, glucose starvation and reactive oxygen species metabolism, together with enrichment of pathways associated with positive regulation of apoptosis and negative regulation of ferroptosis (Fig. 6B). The enrichment of negative regulation of ferroptosis suggests activation of adaptive mechanisms aimed at limiting lipid peroxidation and ferroptotic damage; however, these responses appear insufficient to restore redox homeostasis. Various processes related to cell cycle progression, DNA metabolism and stem cell maintenance were also markedly downregulated, indicating suppression of proliferative and stemness-associated programs following 3-MST inhibition. Enrichment analyses identified significant activation of cell-death-related pathways, including apoptosis, p53 signaling and caspase-associated processes (Supplementary Data 2). Transcriptomic data further indicate the activation of stress and apoptosis-associated programs, with increased expression of key stress regulators such as ATF3, DDIT3 (CHOP) and GADD45B, together with apoptosis-associated genes including BBC3 (PUMA), PMAIP1 (NOXA) and TNFRSF10B (DR5). In parallel, proteomic data indicated activation of antioxidant and stress-adaptive responses, including increased SQSTM1, TXNRD1, SRXN1, SESN2 and AIFM2, consistent with engagement of compensatory survival mechanisms. Together, these data indicate that 3-MST inhibition induces a stress-associated cell-death program integrating apoptosis- and ferroptosis-related pathways.

Transcriptomic analysis showed that changes in protein levels of enzymes involved in central carbon and lipid metabolism closely paralleled changes in the corresponding mRNA levels as WT cancer cells transitioned into the CSC state (Supplementary Data 4). In contrast, this correlation was weak when CSCs were treated with the 3-MST inhibitor, suggesting that alterations in the abundance of key metabolic enzymes in these cells are driven predominantly by post-transcriptional mechanisms or changes in protein stability and degradation (Supplementary Data 4). A similar global pattern emerged when all detected mRNA-protein pairs were compared between WT and CSC cells, where a strong correlation was observed, whereas the correlation was markedly weaker when CSCs were compared with HMPSNE-treated CSCs (Supplementary Data 4). Despite this global divergence between transcript and protein abundance, 3-MST inhibition induced transcriptional activation of several stress-associated pathways (Supplementary Data 1-3). Genes associated with integrated stress signaling and ER stress – including ATF3, DDIT3/CHOP, DDIT4, and GADD45B – were increased. Multiple heat-shock and chaperone genes were also strongly induced. Additional transcriptional changes included regulators of inflammatory signaling and cell death pathways, such as TNFSF13B, TNFSF18, TNFSF4, NFKBIA, and NFKBIZ, as well as genes involved in oxidative stress regulation and iron metabolism, including FTL and SLC7A11, indicating activation of a broad cellular stress response following 3-MST inhibition in CSCs. Transcriptomic and enrichment analyses revealed strong suppression of cell-cycle and DNA replication pathways, with downregulation of key regulators including CDK1, UBE2C and RRM2, consistent with inhibition of proliferative and mitotic programs. Consistently, increased expression of NGFR, SQSTM1, NFKBIA and GADD45B, together with reduced expression of UBE2C and CDK1, further supports loss of proliferative and stemness-associated programs (Supplementary Data 2).

Finally, combined RNA and proteomic analyses revealed suppression of Wnt/β-catenin signaling after 3-MST inhibition in CSCs. WNT16 was reduced at both the transcript and protein levels, while additional components of the pathway, including WLS, TCF7L2, PYGO2 and CCND1, were decreased at the protein level, indicating inhibition of ligand processing, transcriptional activation and downstream output. Hallmark enrichment further confirmed downregulation of Wnt signaling (Supplementary Data 2). Thus, 3-MST inhibition impairs Wnt-signaling, a mechanism essential for CSC maintenance.

## Discussion

The present study identifies 3-MST as an essential metabolic support mechanism for CSCs. CSCs maintain elevated sulfur metabolic activity despite operating in a relatively low-flux metabolic state. Disruption of this pathway compromises CSC proliferation, stemness, and survival, identifying 3-MST as a key metabolic vulnerability.

Consistent with reports that quiescent/dormant CSC-like subpopulations can adopt metabolically restrained states with reduced basal activity [21,22], colon CSCs exhibit reduced oxidative phosphorylation and glycolysis, despite increased expression of sulfur metabolism enzymes and elevated reactive sulfur species. These data suggest that sulfur metabolism supports cellular homeostasis under low bioenergetic conditions, likely through regulation of redox balance and maintenance of mitochondrial function, as reactive sulfur species are known to modulate oxidative stress and electron transport in cancer cells [3–5]. CSCs display increased H_2_S production, reduced ROS levels, enhanced membrane rigidity and marked chemoresistance, linking sulfur metabolism to redox balance and stemness-associated phenotypes. Rescue cells partially reverted these features but retained high 3-MST expression and clonogenic potential, indicating that sustained 3-MST activity represents a persistent component of the CSC state.

MPST deficiency confirmed the functional relevance of this pathway in CSCs. Loss of 3-MST impaired proliferation and migration of colon cancer cells and reduced spheroid formation. Pharmacological inhibition of 3-MST recapitulated the reduction in spheroid growth and induced cell death, while simultaneously causing a marked suppression of cellular bioenergetics. Notably, whereas parental cells retained partial bioenergetic activity after treatment, CSCs, which already operate at a reduced baseline bioenergetic capacity, underwent near-complete suppression of both oxidative phosphorylation and glycolysis. Thus, 3-MST activity is required to sustain minimal metabolic function and survival, and reveal a heightened metabolic vulnerability of CSCs. This finding aligns with prior studies showing that 3-MST supports mitochondrial electron transport and cellular bioenergetics [23–25], maintains glycolysis [26,27], and preserves mitochondrial protein homeostasis [28]. Multi-omics analysis provided mechanistic insights: CSCs show increased abundance of glycolytic and TCA-cycle enzymes despite reduced metabolic flux, consistent with an enzymatically primed but restrained state. Inhibition of 3-MST further destabilizes this architecture by suppressing mitochondrial carbon entry and oxidative metabolism, accompanied by accumulation of pyruvate and lactate, suggesting diversion of carbon away from mitochondrial oxidation. In parallel, proteomic data revealed loss of mitochondrial ribosomal components and respiratory complex I subunits after 3-MST inhibition, indicating impaired mitochondrial protein homeostasis. As mitochondrial respiration depends on continuous complex assembly [29–31], these defects likely drive progressive respiratory failure, which is particularly detrimental in CSCs due to their specific metabolic status.

In addition to metabolic collapse, 3-MST inhibition induced extensive lipid remodeling. Major membrane phospholipids and cholesterol were depleted, whereas free fatty acids, diacylglycerols, and sphingolipid species–including ceramides and sphingosine–accumulated, indicating a shift from structural membrane lipids toward stress-associated lipid species. In parallel, suppression of FADS1 and SCD impaired the generation of protective polyunsaturated and monounsaturated fatty acids. These alterations increased membrane fluidity, consistent with direct biophysical measurements showing reduced membrane rigidity after HMPSNE treatment, and created a lipid environment prone to oxidative damage.

The transcriptional response of CSCs to 3-MST inhibition further reflects a profound cellular stress state, characterized by activation of integrated stress response genes, ER stress markers, heat-shock proteins, and regulators of inflammatory and oxidative stress. These changes are consistent with metabolic and proteotoxic stress triggered by bioenergetic collapse and convergence with mitochondrial dysfunction, redox imbalance, and lipid remodeling to create conditions favorable for iron-dependent lipid peroxidation. Several findings suggest that 3-MST inhibition increases CSC susceptibility to ferroptotic cell death. Proteomic data revealed induction of stress-adaptive pathways related to cystine transport and redox control, including SLC7A11, SLC3A2, PHGDH, PSAT1, ASNS, and TXNRD1, all of which are known to be associated to stem-like phenotypes [32–35]. Simultaneously, key ferroptosis resistance mechanisms were altered: GPX4, a lipid peroxide-reducing enzyme, was reduced, while SAT1 and HMOX1 were strongly induced. HMOX1 promotes labile iron accumulation via heme degradation [36,37], and SAT1 has been implicated in lipid peroxidation and ferroptosis [38,39]. Gene Ontology enrichment further supported activation of oxidative stress, ER stress, glucose starvation, and toxic-response programs, together with pathways linked to positive regulation of apoptosis and negative regulation of ferroptosis, indicating a state of ferroptotic pressure accompanied by compensatory resistance mechanisms. Additional ferroptosis defense systems also appear compromised. Although the ferroptosis suppressor AIFM2/FSP1 was upregulated, proteins involved in coenzyme Q metabolism, such as COQ8B and COQ10A, were reduced, potentially limiting FSP1-CoQ antioxidant capacity [40,41]. Lipid enzyme changes may further heighten vulnerability: reduced stearoyl-CoA desaturase (SCD) may impair monounsaturated fatty acid production that normally buffers peroxidation [42,43], while altered sphingolipid metabolism and increased PEBP1 may promote lipid peroxidation signaling [44]. These alterations indicate that 3-MST inhibition combines induction of ferroptosis-promoting conditions with incomplete compensatory responses, ultimately favoring lipid oxidative damage.

3-MST inhibition also suppressed Wnt/β-catenin signaling, consistent with the known stimulatory effect of H_2_S on this pathway [25,45,46] and its central role in essential CSC functions and viability [14,47,48]. This effect may arise as a downstream consequence of the broader metabolic and stress-related perturbations induced by 3-MST inhibition. The profound bioenergetic collapse would be expected to compromise energy-dependent signaling processes and favor AMPK activation, thereby promoting β-catenin destabilization and reducing TCF7L2-mediated transcription. In parallel, mitochondrial dysfunction and disruption of redox homeostasis, including loss of reactive sulfur species buffering, likely enhance oxidative stress, which is known to impair β-catenin stability and signaling competence. Concomitant lipid remodeling, including reduced cholesterol and increased membrane fluidity, may further disrupt membrane microdomains required for efficient Wnt receptor signaling. Consistently, WNT16 was reduced at both transcript and protein levels, together with decreased abundance of WLS, TCF7L2, PYGO2 and CCND1, indicating suppression of ligand processing, transcriptional activity and consequent proliferation.

Thus, 3-MST inhibition drives CSCs into combined metabolic and structural failure. Mitochondrial dysfunction compromises energy production, while lipid remodeling destabilizes membrane architecture and enhances susceptibility to lipid peroxidation. Concurrent activation of stress responses and impairment of ferroptosis defense mechanisms create conditions permissive for ferroptotic damage. CSC death in response to 3-MST inhibition reflects a mixed stress-associated phenotype integrating ferroptosis, apoptosis and necrosis rather than a single canonical death pathway.

The pronounced lipid remodeling seen in 3-MST-inhibited CSCs suggests an additional layer of destabilization. Lipid and cholesterol biosynthesis are critical dependencies in colorectal CSCs, supporting membrane organization, signaling, and tumor-propagating capacity [43,49]. Altered lipid composition may disrupt membrane-based signaling platforms while increasing membrane fluidity and permeability. These changes may further promote ferroptotic susceptibility, as ferroptosis involves oxidative destruction of membrane phospholipids and is normally counteracted by GPX4 and coenzyme Q systems [41,50]. The combined reduction of GPX4, altered CoQ metabolism, increased SAT1 and PEBP1 expression, and weakened monounsaturated fatty acid buffering likely erode these protective mechanisms and enhance ferroptotic vulnerability. The increase in membrane fluidity, confirmed by functional biophysical measurements, may therefore reflect not only a consequence of lipid remodeling but also destabilization of membrane integrity and signaling.

CSCs are well known for contributing to tumor recurrence and resistance to therapy, yet they are highly refractory to conventional treatments [8–15]. Our results indicate that CSCs depend on 3-MST to maintain mitochondrial stability, bioenergetic balance, and membrane integrity, and targeting this pathway induces global cellular collapse. Notably, 3-MST inhibition suppressed tumor growth in a CSC-derived xenograft model and was effective across multiple colorectal CSC models, supporting its therapeutic relevance. Because 3-MST inhibition simultaneously disrupts multiple essential systems, it may help overcome the metabolic flexibility that allows CSCs to evade targeted therapies and it is fundamentally different from the mode of action of antiproliferative chemotherapy, which is ineffective in CSCs due to their quiescent/dormant nature. The current report identifies 3-MST as a metabolic stabilizer in colorectal CSCs that supports mitochondrial integrity, redox balance, and membrane structure. Inhibiting this pathway disrupts these systems, activates stress responses, and triggers cell death. 3-MST may represent a translatable therapeutic target for selectively eliminating therapy-resistant CSCs.

## Methods

### Cell culture and human specimens

HCT116 human colorectal carcinoma cells (ATCC CCL-247) were obtained from the American Type Culture Collection (ATCC). LoVo human colorectal adenocarcinoma cells (DSMZ ACC-350) were purchased from the German Collection of Microorganisms and Cell Cultures (DSMZ), and SW620 human colorectal adenocarcinoma cells were obtained from the National Cancer Institute (NCI). The epithelial cell line NCM356D, derived from normal colon, was purchased from INCELL Corporation LLC. HCT116 and SW620 cells were cultured in McCoy’s 5A medium (Gibco, cat. no. 26600080), LoVo cells in Ham’s F-12K medium (Gibco, cat. no. 21127022), and NCM356D cells in M3:Base medium (INCELL, cat. no. M300F). All media were supplemented with 10% heat-inactivated fetal bovine serum (FBS; Gibco, cat. no. 10270106) and penicillin–streptomycin (100 U/ml penicillin and 100 μg/ml streptomycin; Gibco, cat. no. 15140122).

Cancer stem cells (CSCs) derived from HCT116, LoVo and SW620 parental cell lines were cultured under serum-free, non-adherent conditions to promote tumorsphere formation. Under these conditions, differentiated cells undergo anoikis, whereas a subpopulation of stem-like, anoikis-resistant cells survives and expands, resulting in enrichment of cancer stem-like cells [51,52]. Subsequent experiments involving CSCs were performed after 3 weeks of selection. Cells were maintained at 37 °C in a humidified incubator with 5% CO_2_. Parental cell lines were passaged twice per week and used for experiments within 2 months after thawing. CSC cultures were passaged once per week, and medium was replaced every 2-3 days.

Fresh-frozen human colon cancer tissues and matched adjacent healthy tissues were obtained from BioIVT (Woodbury, NY, USA). For each patient, tumor tissue and matched non-tumor samples were collected, including adjacent normal tissue (NADJ) or diseased tissue when available. In total, 12 samples were analyzed, comprising six tumor samples and six matched non-tumor samples. The cohort included three male and three female patients. Detailed sample information, including specimen identifiers and tissue classification, is provided in Table 2.

### Cell proliferation and doubling time assays

NCM356D, LoVo and SW620 cells were seeded at 2 × 10⁶ or 4 × 10⁶ cells per T75 flask, and HCT116 cells (wild-type (WT) and MPST-deficient) at 1 × 10⁶ or 2 × 10⁶ cells per T75 flask, in 10 ml of complete culture medium. For CSC enrichment and proliferation assays, 1.8 × 10⁷ HCT116 WT cells were seeded in non-treated T75 flasks in 10 ml of serum-free medium. Under these conditions, differentiated cells do not survive, whereas stem-like cells are selectively enriched and proliferate over time. The high initial seeding density was used to compensate for cell loss during the selection process and the reduced proliferation rate of CSCs under serum-free conditions. Where indicated, MPST-deficient CSCs were subjected to the same enrichment conditions.

Cells were incubated at 37 °C in a humidified atmosphere containing 5% CO_2_ for 72-96 h for NCM356D, LoVo, SW620 and HCT116 cells (WT and MPST-deficient) and up to 10 days for CSCs. At the indicated time points, cells were harvested and viable cell numbers were determined by trypan blue exclusion (1:1 dilution) using LUNA cell counting slides (LUNA, cat. no. L12002) and an automated cell counter (Olympus R1). Doubling time was calculated using nonlinear regression (exponential growth model) in GraphPad Prism based on log-transformed cell counts.

### Western blot analysis

#### Sample preparation

Fresh-frozen colon cancer tissues and matched adjacent healthy tissues were obtained from BioIVT (Woodbury, NY, USA). The cohort included six Caucasian patients (three males and three females, aged 48-75 years). Samples were stored at -80 °C until use. For protein extraction, approximately 30 mg of tissue was transferred into 2 ml microcentrifuge tubes preloaded with three 2.4 mm metal beads and homogenized in 1 ml T-PER™ Tissue Protein Extraction Reagent (Thermo Scientific, cat. no. 78510) supplemented with 1% Halt™ Protease and Phosphatase Inhibitor Cocktail (Thermo Scientific, cat. no. 78445) using a Fisherbrand Bead Mill 4 (one cycle, 120 s, 4 m s⁻¹). Homogenates were centrifuged at 10,000 × g for 20 min at 4 °C, and supernatants containing soluble proteins were collected.

For colon cancer cell lines, NCM356D, LoVo and SW620 cells were seeded at 4 × 10^5^ cells per well, and HCT116 cells at 2 × 10^5^ cells per well, in 6-well plates in 2 ml of complete culture medium. Cells were incubated for 72 h at 37 °C in a humidified atmosphere containing 5% CO_2_ to reach near-confluence, harvested, and lysed as described below.

For HCT116 WT, CSC and Rescue cells, HCT116 WT and Rescue cells were seeded in tissue culture-treated 6-well plates at a density of 2 × 10^5^ cells per well in 2 ml of complete culture medium. HCT116 CSCs were seeded in non-treated 6-well plates at a density of 4.5 × 10^5^ cells per well in 2 ml of serum-free medium. Cells were incubated at 37 °C in a humidified atmosphere containing 5% CO_2_ for 72 h (WT and Rescue) or 8 days (CSCs). For pharmacological treatment, CSCs were treated with HMPSNE (30-300 μM) for 48 h before harvesting. Cells were then collected and lysed as described below.

Cells were scraped from culture plates and lysed in RIPA buffer (Thermo Scientific, cat. no. 89901) supplemented with Halt™ Protease and Phosphatase Inhibitor Cocktail (Thermo Scientific, cat. no. 78445). Lysates were sonicated for 15 s using an ultrasonic bath (XUBA3, Grant Instruments) and clarified by centrifugation. Protein concentration was determined using a bicinchoninic acid (BCA) assay (Thermo Scientific, cat. no. 23225).

### Western blotting

Protein samples were reduced using sample reducing agent (Invitrogen, cat. no. B0009) and denatured in lithium dodecyl sulfate (LDS) sample buffer (Invitrogen, cat. no. B0007) at 95 °C for 5 min. Equal amounts of protein (15-20 μg per lane) were separated by SDS-PAGE on 4-12% Bis-Tris gels (Invitrogen, cat. no. NW04125BOX) and transferred onto PVDF membranes (Invitrogen, cat. no. IB34001X3) using a dry transfer system (iBlot 3, Invitrogen) according to the manufacturer’s instructions. Membranes were blocked in 5% milk (Cell Signaling Technology, cat. no. 9999S) in Tris-buffered saline containing 0.1% Tween-20 (TBST) and incubated with primary antibodies diluted in either 5% BSA (Cell Signaling Technology, cat. no. 9998S) or 5% milk in TBST overnight at 4 °C. Membranes were washed three times for 5 min in TBST and incubated with HRP-conjugated secondary antibodies (1:5,000) for 1 h at room temperature. Signals were detected using chemiluminescence reagents (Radiance Plus, Azure Biosystems) and imaged using an Azure Imaging System 300 (Azure Biosystems). Band intensities were quantified using ImageJ (Fiji) and normalized to β-actin. For human 3-MST, which appears as a doublet corresponding to splice variants, both bands were quantified together. Experimental procedures were performed as previously described with minor modifications [46].

### Reagents and antibodies

The 3-mercaptopyruvate sulfurtransferase (3-MST) inhibitor HMPSNE (2-[(4-hydroxy-6-methylpyrimidin-2-yl)sulfanyl]-1-(naphthalen-1-yl)ethan-1-one) was purchased from MolPort (cat. no. MolPort-000-680-245). The H₂S fluorescent probe 7-azido-4-methylcoumarin (AzMC) was obtained from Sigma-Aldrich (cat. no. 802409). The chemotherapeutic agents paclitaxel (cat. no. HY-B0015) and docetaxel (cat. no. HY-B0011) were purchased from MedChemExpress. Stock solutions were prepared in DMSO and diluted in culture medium to the indicated concentrations. Rabbit monoclonal anti-CBS (D8F2P, cat. no. 14782S), anti-SOD1 (E4G1H, cat. no. 37385S) and HRP-linked anti-mouse IgG secondary antibody (cat. no. 7076S) were purchased from Cell Signaling Technology. Mouse monoclonal anti-β-actin (AC-15, cat. no. A1978) was obtained from Sigma-Aldrich. HRP-conjugated anti-rabbit IgG (H+L) secondary antibody was purchased from Invitrogen (cat. no. 31458). Rabbit polyclonal anti-3-MST (cat. no. ab224043), anti-CSE (cat. no. ab151769), anti-TST (cat. no. ab231248) and rabbit monoclonal anti-ETHE1 (cat. no. ab174302) were obtained from Abcam. Rabbit polyclonal anti-CARS2 (cat. no. HPA041776) and anti-SQRDL (cat. no. HPA017079) were purchased from Sigma-Aldrich.

Mouse monoclonal FITC-conjugated anti-human CD44 (cat. no. 397518), APC-conjugated anti-human CD133 (cat. no. 397906), PE-conjugated anti-human CD166 (cat. no. 343904), APC/Cy7-conjugated anti-human CD29 (cat. no. 303014), Brilliant Violet 711-conjugated anti-human CD326 (EpCAM; cat. no. 324240), PE/Cy5-conjugated anti-human CD26 (cat. no. 302708), Brilliant Violet 605-conjugated anti-human CD24 (cat. no. 311124) and PE/Cy7-conjugated anti-human LGR5 (GPR49; cat. no. 373808) were purchased from BioLegend. Compensation beads (cat. no. 424602) were also obtained from BioLegend.

### Generation of MPST-deficient cell lines with inducible 3-MST re-expression

HCT116 cells with reduced 3-MST expression were generated using a two-step strategy consisting of stable expression of a codon-optimized MPST transgene followed by CRISPR/Cas9-mediated knockout of the endogenous *MPST* gene. A codon-optimized human *MPST* cDNA was synthesized and cloned into a lentiviral inducible expression vector (pLenti-TRE backbone; GenScript). Codon optimization was designed to render the exogenous *MPST* sequence resistant to CRISPR/Cas9-mediated targeting.

Lentiviral particles were produced in HEK293T cells using a third-generation packaging system. Briefly, HEK293T cells were seeded at 1.25 × 10^6^ cells per well in 6-well plates 4-6 h before transfection and transfected with the transfer vector together with packaging plasmids pLP1, pLP2 and pVSV-G using jetOPTIMUS (Polyplus) according to the manufacturer’s instructions. Cells were incubated overnight at 37 °C, after which the medium was replaced, and viral supernatants were collected 24 h later. Supernatants were filtered through a 0.45 μm membrane, aliquoted and stored at -80 °C. Target HCT116 cells were transduced with lentiviral supernatants in the presence of 6 μg/ml protamine sulfate. Seventy-two hours after transduction, cells were subjected to blasticidin selection (12 μg/ml) for 7 days to generate stable populations expressing codon-optimized MPST.

Endogenous *MPST* knockout was subsequently performed in these cells using a lentiviral CRISPR/Cas9 system. A guide RNA targeting human *MPST* (5′-CGTAGGTCTTGATGAAGGCG-3′) was cloned into the pLentiCRISPR v2 vector (GenScript). Lentiviral particles were produced as described above and used to transduce cells already expressing codon-optimized MPST. Seventy-two hours after transduction, cells were selected with puromycin (1 μg/ml) and maintained under selection for 7 days to obtain stable knockout populations. To derive clonal cell lines, cells were plated at low density in 96-well plates and individual colonies were expanded. Knockout of endogenous MPST was confirmed by western blot analysis.

### Tumor-sphere formation assay

HCT116 WT, HCT116 MPST-deficient, HCT116 CSCs, CSC MPST-deficient and Rescue cells were seeded in 48-well plates at a density of 2.5 × 10^4^ cells per well in 25 μL of Matrigel Basement Membrane Matrix (Corning, cat. no. 354234). Plates were incubated at 37 °C in a humidified atmosphere containing 5% CO_2_ for 10 min to allow Matrigel solidification, followed by the addition of 250 μL serum-free McCoy’s 5A medium. Cells were cultured for 8 days at 37 °C and 5% CO_2_, and medium was refreshed every 2-3 days. Spheroids were imaged using an inverted microscope (Olympus CKX53, 10× objective), and spheroid area was quantified using ImageJ (Fiji). For pharmacological experiments, HCT116 cells were treated with HMPSNE (30-300 μM) for 48 h before imaging.

### In vivo tumor growth experiments

All animal experiments were approved by the cantonal veterinary authorities of Fribourg and performed in accordance with Swiss animal welfare regulations (license no. 2023-31-FR). Humanized NSG mice reconstituted with human peripheral blood mononuclear cells (Hu-NSG-PBMC; The Jackson Laboratory) were used for tumor growth experiments. Mice were housed under specific pathogen-free conditions in individually ventilated cages at 21 ± 1 °C on a 12 h light/dark cycle, with ad libitum access to food and water. Animals were acclimatized to the animal facility for 3 weeks prior to experimentation.

For subcutaneous tumor models, mice were injected with 1 × 10^6^ HCT116 WT cells or HCT116-derived CSCs suspended in Matrigel into the flank. Tumor growth was monitored over 25 days by caliper measurements. For comparison of tumor growth between WT- and CSC-derived tumors, experiments were performed in male mice. For pharmacological studies, mice bearing CSC-derived tumors were treated with the 3-MST inhibitor HMPSNE at 3, 10 or 30 mg kg⁻¹ day⁻¹. Treatment was initiated 10 days after tumor cell implantation and administered daily by oral gavage until day 23. Tumor growth was monitored over time using caliper measurements, and tumor volume was calculated as length × width² × 0.5236. Animals were monitored daily for signs of distress and were euthanized before the experimental endpoint if predefined humane criteria were reached. Both male and female mice were included in treatment experiments.

### Apoptosis, necrosis and viability analysis

HCT116 WT and Rescue cells were seeded in tissue culture-treated 96-well plates at a density of 6.7 × 10^3^ cells per well in 100 μL of complete culture medium. HCT116 CSCs, LoVo CSCs and SW620 CSCs were seeded in non-treated 96-well plates at a density of 1.5 × 10^4^ cells per well in 100 μL of serum-free medium. Cells were incubated for 72 h (WT and Rescue) or 8 days (CSCs) at 37 °C in a humidified atmosphere containing 5% CO₂. Treatments with increasing concentrations of docetaxel, paclitaxel or HMPSNE were applied for 48 h before harvesting. Apoptosis and necrosis were assessed using an Annexin V assay as previously described [2]. Briefly, cells were stained with APC-conjugated Annexin V (BioLegend, cat. no. 640920) and propidium iodide (PI; BioLegend, cat. no. 421301) in Annexin V binding buffer (BioLegend, cat. no. 422201) according to the manufacturer’s instructions. Samples were analyzed on a FACS Fortessa flow cytometer (BD Biosciences). APC fluorescence was detected using a 670/30 nm filter (red laser), and PI fluorescence was detected using a 610/20 nm filter (blue laser). Data were analyzed using FlowJo software (v10.10).

### Seahorse extracellular flux analysis

Cellular bioenergetics of HCT116 WT, HCT116 CSCs and Rescue cells were assessed using a Seahorse XFe24 extracellular flux analyzer (Agilent Technologies) as previously described [25]. HCT116 WT and Rescue cells were seeded at 7 × 10^3^ cells per well in 200 μL complete culture medium, and CSCs at 7 × 10^4^ cells per well in 200 μL serum-free medium, in Seahorse XFe24 cell culture microplates. Cells were incubated for 72 h at 37 °C in a humidified atmosphere containing 5% CO₂. For HMPSNE treatment, cells were exposed to increasing concentrations (30-300 μM) for 48 h before the assay.

For mitochondrial respiration measurements (Mito Stress Test), cells were washed twice and incubated in assay medium (XF DMEM, Agilent Technologies, cat. no. 103575-100, pH 7.4) supplemented with 2 mM L-glutamine (Corning, cat. no. 25-015-CL), 1 mM sodium pyruvate (Cytiva, cat. no. SH30239.01) and 10 mM glucose (Gibco, cat. no. A2494001). Plates were equilibrated for 1 h at 37 °C in a CO₂-free incubator. Oxygen consumption rate (OCR) was measured at baseline, followed by sequential injections of oligomycin (1 μM), FCCP (0.2 μM), and rotenone/antimycin A (0.5 μM each) to determine ATP-linked respiration, maximal respiration and non-mitochondrial respiration.

For glycolytic rate measurements, cells were washed and incubated in Seahorse assay medium (XF DMEM, Agilent Technologies, cat. no. 103575-100, pH 7.4) supplemented with 2 mM L-glutamine, 1 mM sodium pyruvate and 10 mM glucose. After 1 h equilibration at 37 °C in a CO₂-free incubator, proton efflux rate (glycoPER) was measured at baseline, followed by sequential injection of rotenone/antimycin A (0.5 μM each) and 2-deoxy-D-glucose (50 mM) to assess compensatory glycolysis and non-glycolytic acidification. Data were analyzed using Wave software (v.2.6, Agilent Technologies) and normalized to total protein content determined by BCA assay (Thermo Scientific, cat. no. 23225).

### Membrane fluidity measurement

Membrane fluidity was assessed using the Membrane Fluidity Assay Kit (Abcam, cat. no. ab189819). HCT116 WT and Rescue cells were seeded in tissue culture-treated 96-well plates at a density of 6.7 × 10^3^ cells per well in 100 μL of complete culture medium, whereas HCT116 CSCs were seeded in non-treated 96-well plates at a density of 1.5 × 10^4^ cells per well in 100 μL of serum-free medium. Cells were plated in triplicate and incubated for 72 h (WT and Rescue) or 8 days (CSCs) at 37 °C in a humidified atmosphere containing 5% CO_2_. Where indicated, cells were treated with HMPSNE (30-300 μM) for 48 h before the endpoint. For membrane labeling, a solution containing 5 μM fluorescent lipid reagent in perfusion buffer was prepared and supplemented with Pluronic F127 to a final concentration of 0.08% (from a 1% stock solution). Cells were incubated with the labeling solution for 1 h at 25 °C in the dark. Plates were then centrifuged at 2,500 rpm for 1 min, and cells were washed twice with culture medium to remove unincorporated dye. Cells were resuspended in fresh medium, and fluorescence was measured using a SpectraMax iD5 plate reader with excitation at 350 nm and emission at 400 nm (monomer) and 470 nm (excimer). Background fluorescence was subtracted, and the excimer-to-monomer fluorescence ratio (470/400 nm) was calculated.

### Reactive oxygen species measurement

Intracellular reactive oxygen species (ROS) levels were measured using the DCFDA/H₂DCFDA Cellular ROS Assay Kit (Abcam, cat. no. ab113851). HCT116 WT and Rescue cells were seeded in black, clear-bottom tissue culture-treated 96-well plates at a density of 6.7 × 10^3^ cells per well in 100 μL of complete medium, and HCT116 CSCs were seeded in non-treated 96-well plates at a density of 1.5 × 10^4^ cells per well in 100 μL of serum-free medium. Cells were plated in triplicate and incubated for 72 h (WT and Rescue) or 8 days (CSCs) at 37 °C in a humidified atmosphere containing 5% CO₂. For HMPSNE treatment, cells were exposed to increasing concentrations (30-300 μM) for 48 h before the assay. Cells were washed and incubated with 20 μM DCFDA in assay buffer for 45 min at 37 °C in the dark. Following incubation, cells were washed and fluorescence was measured immediately using a SpectraMax iD5 microplate reader (Molecular Devices) at excitation/emission wavelengths of 485/535 nm. Fluorescence values were normalized to total protein content determined using a BCA assay (Thermo Scientific, cat. no. 23225).

### Measurement of intracellular polysulfides (H_2_S_n_)

Intracellular polysulfide levels were measured using the SSP4 fluorescent probe (Dojindo Molecular Technologies, cat. no. SB10). HCT116 WT and Rescue cells were seeded in black, clear-bottom tissue culture-treated 96-well plates at a density of 6.7 × 10^3^ cells per well in 100 μL of complete culture medium, whereas HCT116 CSCs were seeded in non-treated 96-well plates at a density of 1.5 × 10^4^ cells per well in 100 μL of serum-free medium. Cells were plated in triplicate and incubated for 72 h (WT and Rescue) or 8 days (CSCs) at 37 °C in a humidified atmosphere containing 5% CO₂. Following incubation, cells were washed with serum-free medium and incubated with 10 μM SSP4 in serum-free medium supplemented with 0.5 mM cetyltrimethylammonium bromide (CTAB) for 15 min at 37 °C. Cells were then washed twice with PBS, and fluorescence was measured in PBS using a SpectraMax iD5 plate reader (Molecular Devices) at excitation/emission wavelengths of 482/515 nm. Fluorescence values were normalized to total protein content determined using a BCA assay (Thermo Scientific, cat. no. 23225).

### Flow cytometry analysis of CSC markers

HCT116 WT cells were seeded in tissue culture-treated 96-well plates at a density of 6.7 × 10^3^ cells per well in 100 μL of complete medium, and HCT116 CSCs were seeded in non-treated 96-well plates at a density of 1.5 × 10^4^ cells per well in 100 μL of serum-free medium. WT cells and CSCs were incubated for 72 h and 8 days, respectively, at 37 °C in a humidified atmosphere containing 5% CO₂. Where indicated, cells were treated with increasing concentrations of HMPSNE for 48 h before analysis. Cells were harvested, washed with FACS buffer, and incubated with fluorophore-conjugated antibodies against CD44 (FITC), CD133 (APC), CD166 (PE), CD29 (APC/Cy7), EpCAM (CD326; BV711), CD26 (PE/Cy5), CD24 (BV605) and LGR5 (PE/Cy7) for 15 min at 4 °C in the dark. Compensation beads (BioLegend, cat. no. 424602) were used for multicolor compensation. Samples were analyzed using a BD LSRFortessa flow cytometer (BD Biosciences), and data were processed using FlowJo software (v10.10.0).

### H_2_S detection using AzMC

Intracellular H_2_S levels were measured using the fluorescent probe 7-azido-4-methylcoumarin (AzMC) as previously described [53]. HCT116 WT and Rescue cells were seeded in black, clear-bottom tissue culture-treated 96-well plates at a density of 6.7 × 10^3^ cells per well in 100 μL of complete medium, and HCT116 CSCs were seeded in non-treated 96-well plates at a density of 1.5 × 10^4^ cells per well in 100 μL of serum-free medium. WT and Rescue cells were incubated for 72 h, and CSCs for 8 days, at 37 °C in a humidified atmosphere containing 5% CO₂. For HMPSNE treatment, cells were exposed to increasing concentrations (30-300 μM) for 48 h before the assay. Cells were washed twice with HBSS (Gibco, cat. no. 14025050) and incubated for 1 h at 37 °C in HBSS containing 10 μM AzMC. Fluorescence imaging was performed using a Cytation 5 imaging reader (Agilent) with 10× and 20× objectives. Because of the spheroid morphology of CSCs, fluorescence intensity was quantified using a SpectraMax iD5 microplate reader (Molecular Devices) for all cell types to ensure consistency across conditions, and values were normalized to total protein content. For pharmacological experiments, CSCs were treated with increasing concentrations of HMPSNE (30-300 μM) for 48 h before AzMC incubation.

### ’Wound healing’ cell migration assay

HCT116 WT, HCT116 MPST-deficient, HCT116 CSCs and CSC MPST-deficient cells were seeded in 96-well plates at a density of 5 × 10^4^ cells per well in 100 μL of complete culture medium and incubated for 24 h at 37 °C in a humidified atmosphere containing 5% CO_2_ to allow formation of a confluent monolayer. Uniform wounds were generated using the WoundMaker (Essen BioScience), after which the medium was replaced with fresh complete culture medium to remove detached cells. Plates were transferred to a Cytation 5 imaging system (Agilent BioTek) equipped with a 10× objective, and phase-contrast images were acquired every 4 h for 48 h under standard culture conditions (37 °C, 5% CO_2_). Cell migration was quantified by measuring wound confluence over time using Gen5 software.

### Telomere length measurement

Genomic DNA was extracted using the NucleoSpin DNA RapidLyse kit (Macherey-Nagel) according to the manufacturer’s instructions. Briefly, up to 1 × 10^6^ cells were lysed in lysis buffer containing Proteinase K and incubated at 56 °C until complete lysis, followed by binding of DNA to a silica membrane, washing, and elution in 100 μL elution buffer. DNA concentration and purity were determined using a NanoDrop spectrophotometer (Thermo Fisher Scientific). Telomere length was quantified using the Absolute Human Telomere Length Quantification qPCR Assay Kit (ScienCell, cat. no. 8918) according to the manufacturer’s instructions. For each sample, quantitative PCR reactions were performed in parallel using telomere-specific primers and a single-copy reference (SCR) primer set. Reactions (20 μL) contained genomic DNA (5 ng), primer mix, and 2× SYBR Green master mix. qPCR was performed using the following cycling conditions: initial denaturation at 95 °C for 10 min, followed by 32 cycles of 95 °C for 20 s, 52 °C for 20 s, and 72 °C for 45 s, with data acquisition at each cycle. Relative telomere length was calculated using the comparative ΔΔCq method by normalizing telomere amplification to the single-copy reference gene and to a reference genomic DNA sample with known telomere length provided in the kit.

### OMICS

#### RNA sequencing and data analysis

HCT116 WT and Rescue cells were seeded in tissue culture-treated 6-well plates at a density of 2 × 10^5^ cells per well in 2 ml of complete medium. HCT116 cancer stem cells (CSCs) were seeded in non-treated 6-well plates at a density of 4.5 × 10^5^ cells per well in 2 ml of serum-free medium. Cells were incubated at 37 °C in a humidified atmosphere with 5% CO_2_ for 72 h (WT and Rescue) or 8 days (CSCs). For pharmacological treatment, cells were treated with 300 μM HMPSNE for 48 h prior to harvesting. Cells were collected and stored at -80 °C until RNA extraction. Total RNA was extracted using the NucleoSpin® RNA Plus kit (Macherey-Nagel, cat. no. 740984) according to the manufacturer’s instructions. RNA concentration and purity were assessed using a NanoDrop spectrophotometer (Thermo Fisher Scientific). RNA quality control, including RNA integrity assessment using a Fragment Analyzer, as well as library preparation and sequencing, were performed at the Genomic Technologies Facility (University of Lausanne, Switzerland). RNA-seq libraries were prepared using a stranded mRNA library preparation protocol and sequenced on an Aviti platform (Element Biosciences). Single-end sequencing (150 bp) was performed, with an average depth of ∼25 million reads per sample.

Sequencing reads were processed using a standard pipeline including adapter trimming, quality filtering, and alignment to the human reference genome (GRCh38). Gene-level count matrices were generated by the sequencing facility and used for downstream analysis. Five biological replicates per condition were analyzed.

Differential expression analysis was performed in R (v4.5.2) using DESeq2. Genes with very low expression were filtered by retaining those with a total read count ≥10 across all samples. Raw counts were normalized using the median-of-ratios method. Differential expression was assessed using a negative binomial generalized linear model with design condition. Comparisons were performed between CSCs_CTR vs WT and CSCs_HMPSNE vs CSCs_CTR. Log2 fold changes were shrunk using the apeglm method. Genes with false discovery rate (FDR) < 0.05 and absolute fold change ≥1.5 were considered significant. For visualization, variance stabilizing transformation (VST) was applied to normalized counts. Principal component analysis (PCA) was performed to assess global transcriptomic differences between conditions, revealing clear separation between groups. Volcano plots were generated using shrunken log2 fold changes, and the top 15 upregulated and downregulated genes were annotated. Heatmaps were generated from the 50 most variable genes using row-scaled (z-score) VST-transformed counts.

Additional exploratory analyses were performed using iDEP 2.0 and are presented in the supplementary data, including clustering and additional visualizations. Genes with low expression were filtered using a threshold of 1 CPM in at least one sample, and data were transformed using log2(CPM + 1). Results obtained using iDEP were consistent with DESeq2 analysis. Functional enrichment analyses were performed using gene set enrichment analysis (GSEA) and over-representation analysis (ORA). GSEA was conducted using WebGestalt with Homo sapiens as the reference organism and Ensembl gene identifiers as input. Gene sets from Gene Ontology (GO) Biological Process and Cellular Component, KEGG, Reactome, and Hallmark collections were analyzed using 1,000 permutations, with gene set sizes ranging from 5 to 2000 genes. Categories with FDR < 0.05 were considered significantly enriched. Over-representation analysis (ORA) was performed using Metascape. Gene identifiers were mapped to Entrez gene IDs prior to analysis. Enrichment analysis was conducted using Gene Ontology (GO) Biological Process and Cellular Component, KEGG, Reactome, and Hallmark gene sets, with genes detected in the RNA-seq dataset used as background. Enrichment significance was assessed using a cumulative hypergeometric test with Benjamini–Hochberg correction. Terms with p-value < 0.05, enrichment factor > 1.5, and a minimum of 3 genes were considered significant. Enriched terms were grouped into clusters based on similarity using kappa statistics.

### Proteomics data processing and analysis

HCT116 WT and Rescue cells were seeded in tissue culture-treated 6-well plates at a density of 2 × 10^5^ cells per well in 2 ml of complete culture medium. HCT116 CSCs were seeded in non-treated 6-well plates at a density of 4.5 × 10^5^ cells per well in 2 ml of serum-free medium. Cells were incubated at 37 °C in a humidified atmosphere containing 5% CO_2_ for 72 h (WT and Rescue) or 8 days (CSCs). For pharmacological treatment, cells were treated with HMPSNE 300 μM for 48 h before harvesting. Cells were then collected and frozen at -80°C until protein extraction from Metabolomics and Proteomics Platform (MAPP) at UNIFR. Proteins were extracted in 1% sodium deoxycholate, reduced with dithiothreitol and alkylated with iodoacetamide, followed by overnight digestion with trypsin (1:50, enzyme:protein). Peptides were acidified, clarified by centrifugation, and desalted using StageTips. LC–MS/MS analysis was performed on an Exploris 480 mass spectrometer (Thermo Scientific) coupled to a Vanquish NEO UHPLC system. Peptides were separated on a 75 μm × 20 cm C18 column using a linear gradient of acetonitrile in 0.1% formic acid. Data were acquired in data-independent acquisition (DIA) mode over a mass range of m/z 350–1200. Raw data were processed using Spectronaut (v18, Biognosys) with the directDIA+ workflow against the UniProt human database including common contaminants. Downstream statistical analysis was performed in Perseus using permutation-based false discovery rate (FDR < 0.05). Five biological replicates per condition were analyzed. Proteomics intensity values were log2-transformed prior to statistical analysis. Missing values were imputed by the proteomics facility using Perseus software prior to downstream analysis. Data were analyzed in R (v4.5.2). Proteins exhibiting zero variance across samples were removed prior to analysis. Differential protein abundance between experimental groups was assessed using the limma package, which applies linear models with empirical Bayes moderation. A design matrix was constructed to model the three experimental conditions (WT, CSCs_CTR, and CSCs_HMPSNE), and pairwise contrasts were defined to compare CSCs_CTR vs WT and CSCs_HMPSNE vs CSCs_CTR. Resulting p-values were adjusted for multiple testing using the Benjamini–Hochberg false discovery rate (FDR) method. Proteins with FDR < 0.05 and absolute fold change ≥ 1.5 were considered significantly altered. Principal component analysis (PCA) was performed using centered and scaled protein intensity values. Volcano plots were generated to visualize the relationship between log2 fold change and statistical significance (−log10 FDR), and the top 15 upregulated and downregulated proteins were annotated. Heatmaps were generated from the 50 most variable proteins across samples based on variance. Protein intensities were scaled by row (z-score), with values capped between −2 and 2 to improve visualization. Samples were displayed in biological order without clustering. The visualizations were performed in R using the packages limma, ggplot2, and pheatmap.

Additional analyses were performed using MetaboAnalyst (version 6.0) and are presented in the supplementary data. Data were normalized by median, log2-transformed, and autoscaled prior to analysis. No filtering was applied. Principal component analysis (PCA), volcano and heatmaps were generated to assess global patterns in the data. Statistical significance of group separation was evaluated using permutational multivariate analysis of variance (PERMANOVA) with 999 permutations. Differential features were identified based on fold change (≥1.5) and p-value < 0.05.

Functional enrichment analyses were performed using gene set enrichment analysis (GSEA) and over-representation analysis (ORA). For proteomic datasets, gene identifiers (or corresponding gene symbols derived from protein identifiers) were used as input. GSEA was conducted using WebGestalt with Homo sapiens as the reference organism. Gene sets from Gene Ontology (GO) Biological Process and Cellular Component, KEGG, Reactome, and Hallmark collections were analyzed using 1,000 permutations, with gene set sizes ranging from 5 to 2000 genes. Categories with a false discovery rate (FDR) < 0.05 were considered significantly enriched. Over-representation analysis (ORA) was performed using Metascape. Gene identifiers were mapped to Entrez gene IDs prior to analysis. Enrichment analysis was conducted using Gene Ontology (GO) Biological Process and Cellular Component, KEGG, Reactome, and Hallmark gene sets, with genes detected in the corresponding dataset used as background. Enrichment significance was assessed using a cumulative hypergeometric test with Benjamini–Hochberg correction. Terms with p-value < 0.05, enrichment factor > 1.5, and a minimum of 3 genes were considered significant. Enriched terms were grouped into clusters based on similarity using kappa statistics. To reduce redundancy and facilitate interpretation, REVIGO analysis was performed on significantly enriched GO Biological Process terms using a dispensability threshold of 0.5. For visualization purposes, GO term lists derived from exploratory analyses using MetaboAnalyst were used, yielding results consistent with the primary R-based analysis.

#### Metabolomics

HCT116 WT and Rescue cells were seeded in tissue culture-treated 10 cm² dishes at a density of 1.6 × 10^6^ cells per dish in 10 ml of complete medium. HCT116 cancer stem cells (CSCs) were seeded in non-treated 10 cm² dishes at a density of 3.5 × 10^6^ cells per dish in 10 ml of serum-free medium. Cells were incubated at 37 °C in a humidified atmosphere with 5% CO_2_ for 72 h (WT and Rescue) or 8 days (CSCs). For pharmacological treatment, cells were treated with 300 μM HMPSNE for 48 h prior to harvesting. Cells were washed twice with PBS and submitted to the Metabolomics & Lipidomics Facility (University of Lausanne, Switzerland) for analysis. Polar metabolites were extracted using ice-cold 80% methanol, followed by mechanical homogenization and centrifugation. Supernatants were dried and reconstituted in 80% methanol, with volumes normalized to total protein content determined by BCA assay. Targeted metabolomics analysis was performed by hydrophilic interaction liquid chromatography coupled to tandem mass spectrometry (HILIC–MS/MS) using a 6495 triple quadrupole mass spectrometer interfaced with a 1290 UHPLC system (Agilent Technologies). Data were acquired in dynamic multiple reaction monitoring (dMRM) mode, enabling the detection of over 400 metabolites involved in central carbon metabolism and related pathways.

Raw data were processed using MassHunter software (Agilent Technologies). Relative metabolite abundances were determined from extracted ion chromatogram peak areas. Signal drift was corrected using a locally weighted scatterplot smoothing (LOWESS) approach based on pooled quality control (QC) samples analyzed throughout the batch. Metabolites with high analytical variability (coefficient of variation >30% in QC samples) were excluded from further analysis. Five biological replicates per condition were analyzed.

Metabolomics data were analyzed in R (v4.5.2) using the limma package. Input data consisted of metabolite abundances expressed as peak area normalized to protein content. Metabolite names were curated to remove empty entries and ensure unique identifiers. Raw values were log2-transformed after addition of a pseudocount defined as half of the smallest positive value in the dataset. Metabolites with non-finite values or zero variance across samples were removed prior to analysis. A design matrix was constructed to model the three experimental conditions (WT, CSCs_CTR, and CSCs_HMPSNE). Differential metabolite abundance was assessed using linear models with empirical Bayes moderation implemented in limma. Pairwise contrasts were defined to compare CSCs_CTR vs WT and CSCs_HMPSNE vs CSCs_CTR. Resulting p-values were adjusted for multiple testing using the Benjamini–Hochberg false discovery rate (FDR) method. Metabolites with FDR < 0.05 and absolute fold change ≥ 1.5 were considered significantly altered. Principal component analysis (PCA) was performed on log2-transformed metabolite intensities after centering and scaling. Volcano plots were generated using log2 fold change and −log10(FDR), and the top 15 upregulated and downregulated metabolites were annotated. Heatmaps were generated from the 50 most variable metabolites across samples based on variance. Metabolite intensities were scaled by row (z-score), with values capped between −2 and 2 to improve visualization. Samples were displayed in biological order without clustering. The visualizations were performed in R using the packages limma, ggplot2, and pheatmap.

Additional analyses were performed using MetaboAnalyst (version 6.0) and are presented in the supplementary data. Data were imported as peak intensity values (CSV format) with samples in columns and analyzed as unpaired data. Metabolite abundances were pre-normalized to protein content (peak area per µg of protein) prior to analysis; therefore, no additional sample normalization was applied in MetaboAnalyst. Data were log10-transformed to reduce heteroscedasticity and approximate normality, followed by autoscaling (mean-centering and division by the standard deviation of each variable). Low-variance filtering was applied using the interquartile range (IQR) method, removing the lowest 5% of variables to reduce noise. Statistical analysis was performed using one-way analysis of variance (ANOVA). Metabolites with p-value < 0.05 and fold change ≥ 1.5 were considered significantly altered. Principal component analysis (PCA) and hierarchical clustering heatmaps were generated to assess global metabolic differences between conditions. Statistical significance of group separation was evaluated using permutational multivariate analysis of variance (PERMANOVA) with 999 permutations. Functional enrichment analyses were performed using both quantitative enrichment analysis (QEA) and over-representation analysis (ORA) implemented in MetaboAnalyst. QEA was conducted using all detected metabolites to identify coordinated pathway-level changes based on quantitative differences across conditions. ORA was performed using significantly altered metabolites to identify over-represented metabolic pathways, with the list of metabolites detected in the dataset used as background. Enrichment analyses were conducted using curated metabolic pathway databases integrated within MetaboAnalyst (including KEGG-based metabolite sets). Pathways with p-value < 0.05 were considered significantly enriched.

#### Lipidomics

HCT116 WT and Rescue cells were seeded in tissue culture-treated 10 cm² dishes at a density of 1.6 × 10^6^ cells per dish in 10 ml of complete medium. HCT116 cancer stem cells (CSCs) were seeded in non-treated 10 cm² dishes at a density of 3.5 × 10^6^ cells per dish in 10 ml of serum-free medium. Cells were incubated at 37 °C in a humidified atmosphere with 5% CO₂ for 72 h (WT and Rescue) or 8 days (CSCs). For pharmacological treatment, cells were treated with 300 μM HMPSNE for 48 h prior to harvesting. Cells were washed twice with PBS and submitted to the Metabolomics & Lipidomics Facility (University of Lausanne, Switzerland) for analysis. Lipids were extracted from cells using ice-cold 80% methanol, followed by homogenization and centrifugation. Supernatants were dried and reconstituted in isopropanol containing a mixture of isotopically labeled internal standards covering multiple lipid classes. Lipidomic analysis was performed using hydrophilic interaction liquid chromatography coupled to electrospray ionization tandem mass spectrometry (HILIC–ESI–MS/MS) on a TSQ Altis triple quadrupole mass spectrometer (Thermo Fisher Scientific). Data were acquired in timed selected reaction monitoring (tSRM) mode in both positive and negative ionization modes, enabling the quantification of approximately 780 lipid species across major lipid classes, including glycerolipids, glycerophospholipids, sphingolipids, cholesterol esters, and free fatty acids. Raw data were processed using TraceFinder software (Thermo Fisher Scientific). Lipid abundances were estimated based on internal standard normalization to account for differences in ionization and fragmentation efficiency. Five biological replicates per condition were analyzed.

Lipidomics data were analyzed in R (v4.5.2) using the limma package. Input data consisted of lipid abundances expressed as mol% of total lipid content. Lipid names were curated to remove empty entries and ensure unique identifiers. Raw values were log2-transformed after addition of a pseudocount defined as half of the smallest positive value in the dataset. Lipids were retained for differential analysis if they were detected in at least three replicates in at least one experimental group. Lipids with non-finite values or zero variance after transformation were removed prior to analysis. A design matrix was constructed to model the three experimental conditions (WT, CSCs_CTR, and CSCs_HMPSNE). Differential lipid abundance was assessed using linear models with empirical Bayes moderation implemented in limma. Pairwise contrasts were defined to compare CSCs_CTR vs WT and CSCs_HMPSNE vs CSCs_CTR. Resulting p-values were adjusted for multiple testing using the Benjamini–Hochberg false discovery rate (FDR) method. Lipids with FDR < 0.05 and absolute fold change ≥ 1.5 were considered significantly altered. Principal component analysis (PCA) was performed on centered and scaled log2-transformed data. Heatmaps were generated using the top 50 most variable lipids, scaled by row (z-score) and capped between −2 and 2. Volcano plots were generated with the most significantly altered features were ranked based on log2 fold change, and the top 15 upregulated and top 15 downregulated lipids were visualized. Lipid class composition was calculated by summing mol% values per lipid class. To investigate lipid structural properties, fatty acyl chain length and degree of unsaturation were extracted from lipid annotations. Lipid distributions were visualized using chain length × unsaturation heatmaps and corresponding fold-change maps. The unsaturation index (UI) was calculated as the mol%-weighted sum of double bonds, and average chain length (ACL) as the mol%-weighted average carbon number.

Additional analyses were performed using MetaboAnalyst (version 6.0) and are presented in the supplementary data. Data were used as provided by the analytical platform (mol% of total lipid content), without additional normalization. Data were log-transformed (base 10) and autoscaled prior to analysis. PCA, heatmaps, and volcano plots were generated, and top 25 increased and decreased lipid features were identified. Multivariate statistical significance was assessed using PERMANOVA with 999 permutations based on Euclidean distances. Over-representation analysis (ORA) was performed in MetaboAnalyst using significantly altered lipids to identify enriched lipid-related pathways or classes. The set of lipids detected in the dataset was used as background. Lipid ontology enrichment analysis was performed using the LION web tool in ranking mode and target-list mode. Analyses were performed separately for increased and decreased lipid subsets where applicable. Significant ontology terms were defined at FDR < 0.05. Enrichment results are presented in the supplementary data, with selected terms shown in Figure 6. These results were visualized in R as bubble plots, where the x-axis represents ontology terms, the y-axis corresponds to enrichment significance (−log10(FDR)), point size indicates the number of annotated lipids, and color reflects enrichment significance.

### Statistical analysis

Data are presented as mean ± SEM unless otherwise indicated. Statistical analyses were performed using GraphPad Prism (GraphPad Software). For comparisons between two groups, unpaired two-tailed t-tests were used, with Welch’s correction applied when variances were unequal. For comparisons of more than two groups, one-way ANOVA followed by Dunnett’s or Tukey’s multiple comparisons test was performed as appropriate. For experiments involving two independent variables, two-way ANOVA followed by Sidak’s multiple comparisons test was used. When data did not meet assumptions of normality, non-parametric tests were applied, including the Kruskal-Wallis test followed by Dunn’s multiple comparisons test. Differences were considered statistically significant at *P* < 0.05.

## Data Availability

All raw data and source data are available from the corresponding authors upon reasonable request. Source data for Western blot analyses are available at Zenodo (https://doi.org/10.5281/zenodo.19221600). RNA-seq data have been submitted to GEO/SRA under accession SUB16081162 and will be released upon publication. Proteomics data will be deposited in the PRIDE repository, and metabolomics and lipidomics data will be deposited in the MetaboLights repository. These datasets are currently under processing and accession numbers will be provided upon completion.

## Acknowledgements

We thank Dr. Anita Marton, Maeva Oggier, Dr. Jimmy Stalin, Oriana Coquoz and Dr. Theodora Panagaki for technical support and Dr. Milos Filipovic for helpful suggestions regarding omics data analysis and presentation. We also acknowledge the following facilities: at the University of Fribourg, the Cell Analytics Facility (CAF), the Metabolomics and Proteomics Platform (MAPP) and the Bioinformatics & Biostatistics Core Facility; and at the University of Lausanne, the Genomic Technologies Facility (GTF) and the Metabolomics & Lipidomics Facility. This work was supported by the University of Fribourg Research Pool (Grant #PL-25-07) and the University of Fribourg Centenary Research Fund (Grant #FC-24-932) (to K.A.), the Swiss Krebsliga (Grant #KLS 4504-08-2018R) and the Novartis Foundation (Grant #23B110) (to C.S.).

## Author Contributions Statement

Conceptualization: C.S. and K.A.; Methodology, experimentation, and data analysis: K.A. and O.O.; Funding acquisition: C.S. and K.A.; Project administration: K.A. and C.S.; Supervision: K.A.; Writing-original draft: C.S. and K.A.; Writing-review and editing: K.A. and C.S.

## Competing Interests

The authors declare no competing interests.

## Correspondence and Requests for Materials

should be addressed to Kelly Ascenção or Csaba Szabo.

**Supplementary Data 1 | Integrated multi-omics profiling of colorectal cancer stem cells (CSCs) compared with HCT116 wild-type (WT) cells.** Comprehensive transcriptomic, proteomic, metabolomic and lipidomic analyses comparing CSCs and WT cells. The dataset includes principal component analysis (PCA) to assess sample clustering and separation between conditions, hierarchical clustering and heatmaps of the most variable or differentially regulated features, and differential expression analyses visualized by volcano plots. Lists of the top 25 upregulated and downregulated genes, proteins, metabolites and lipid species are also provided. For transcriptomic datasets, differential expression analyses were performed using fold-change and statistical significance thresholds (fold change ≥1.5 and false discovery rate (FDR) ≤0.05). For proteomics, metabolomic and lipidomic analyses, significantly altered metabolites and lipid species were identified using fold change ≥1.5 and P value ≤0.05. PCA plots include PERMANOVA statistics to evaluate group separation. Functional interpretation of differential features was performed using pathway and enrichment analyses appropriate for each omics layer. For transcriptomic and proteomic datasets, Gene Set Enrichment Analysis (GSEA) was performed using Gene Ontology (GO) biological process and cellular component annotations, as well as KEGG, Reactome and Hallmark gene sets. In addition, Over-Representation Analysis (ORA) was used to identify biological processes and pathways significantly enriched among genes or proteins increased or decreased in CSCs relative to WT cells. For metabolomic datasets, pathway enrichment was assessed using Quantitative Enrichment Analysis (QEA), and ORA was used to identify metabolite sets or pathways enriched among metabolites altered between conditions. For lipidomic datasets, ORA was performed to identify enriched lipid-associated pathways or annotations among lipid species altered between conditions. Lipid ontology enrichment was further evaluated using the LION (Lipid Ontology) framework in both ranking and target-list modes to identify structural and functional lipid categories associated with CSCs or WT cells. Together, these analyses provide an integrated multi-omics characterization of molecular differences between colorectal cancer stem cells and their parental HCT116 wild-type counterparts.

**Supplementary Data 2 | Multi-omics profiling of pharmacological 3-MST inhibition in colorectal cancer stem cells.** Integrated transcriptomic, proteomic, metabolomic and lipidomic analyses comparing cancer stem cells treated with the 3-MST inhibitor HMPSNE and untreated CSC controls (CSCs_CTR). The dataset includes principal component analysis (PCA), hierarchical clustering, heatmaps of the most variable or differentially regulated features, volcano plots of differential expression, and lists of the top 25 upregulated and downregulated genes, proteins, metabolites and lipid species. Differential expression analyses were performed using a fold-change threshold ≥1.5 with statistical significance defined as false discovery rate (FDR) ≤0.05 for transcriptomic datasets and P ≤0.05 for proteomic, metabolomic and lipidomic datasets. Functional interpretation of differential features was performed using pathway and enrichment analyses appropriate for each omics layer. For transcriptomic and proteomic datasets, Gene Set Enrichment Analysis (GSEA) was performed using Gene Ontology (GO) biological process and cellular component annotations as well as KEGG, Reactome and Hallmark gene sets. Over-Representation Analysis (ORA) was additionally used to identify pathways and biological processes significantly enriched among features increased or decreased following HMPSNE treatment. For metabolomic datasets, pathway enrichment was assessed using Quantitative Enrichment Analysis (QEA) to evaluate coordinated changes in metabolite sets across metabolic pathways. ORA was additionally applied to identify metabolite pathways significantly associated with HMPSNE treatment. For lipidomic datasets, enrichment of lipid pathways and lipid-associated annotations was assessed using ORA. Lipid ontology enrichment was further evaluated using the LION (Lipid Ontology) framework in both ranking and target-list modes to identify structural and functional lipid categories altered following 3-MST inhibition. The dataset additionally includes integrated multi-omics comparisons across three experimental conditions (WT, CSCs_CTR and CSCs_HMPSNE), including PCA analyses and differential profiles across transcriptomic, proteomic, metabolomic and lipidomic datasets. Lipid structural properties, including fatty-acid chain length and unsaturation indices, were also analyzed to characterize lipid remodeling associated with CSC state and 3-MST inhibition. Together, these analyses provide a comprehensive multi-omics characterization of molecular changes induced by pharmacological inhibition of 3-MST in colorectal cancer stem cells.

**Supplementary Data 3 | Multi-omics profiling of pharmacological 3-MST inhibition in HCT116 wild-type colorectal cancer cells.** Integrated transcriptomic, proteomic, metabolomic and lipidomic analyses comparing HCT116 wild-type (WT) cells treated with the 3-mercaptopyruvate sulfurtransferase (3-MST) inhibitor HMPSNE and untreated WT control cells (WT_CTR). The dataset includes principal component analysis (PCA), hierarchical clustering, heatmaps of the most variable or differentially regulated features, volcano plots of differential expression, and lists of the top 25 upregulated and downregulated genes, proteins, metabolites and lipid species. Differential expression analyses were performed using a fold-change threshold ≥1.5 with statistical significance defined as false discovery rate (FDR) ≤0.05 for transcriptomic datasets and P ≤0.05 for proteomic, metabolomic and lipidomic datasets. Functional interpretation of differential features was performed using pathway enrichment analyses. For transcriptomic and proteomic datasets, Gene Set Enrichment Analysis (GSEA) was conducted using Gene Ontology (GO) biological process and cellular component annotations, as well as KEGG, Reactome and Hallmark gene sets. Over-Representation Analysis (ORA) was additionally performed to identify pathways and biological processes significantly enriched among features increased or decreased following HMPSNE treatment. For metabolomic datasets, pathway enrichment was evaluated using Quantitative Enrichment Analysis (QEA) to identify metabolic pathways significantly altered following pharmacological inhibition of 3-MST. ORA was also applied to determine metabolite pathways significantly associated with treatment. For lipidomic datasets, enrichment of lipid pathways was assessed using ORA. Lipid ontology enrichment was further evaluated using the LION (Lipid Ontology) framework in both ranking and target-list modes to identify lipid structural and functional categories associated with HMPSNE treatment. Together, these analyses provide a comprehensive multi-omics characterization of molecular changes induced by pharmacological inhibition of 3-MST in HCT116 wild-type colorectal cancer cells.

**Supplementary Data 4 | Correlation between transcriptomic and proteomic changes in WT HCT116 cells vs. CSCs and in CSCs after HMPSNE treatment.** Scatter plots showing the relationship between RNA expression changes (RNA log_2_ fold change) and protein abundance changes (protein log_2_ fold change) derived from integrated transcriptomic and proteomic datasets. Top panels: Correlation analysis restricted to metabolic enzymes. The left panel compares CSCs with WT cells, showing a moderate correlation between RNA and protein changes (Pearson r = 0.65). The right panel compares HMPSNE-treated CSCs with untreated CSCs (CSC+HMPSNE vs CSC), showing a weaker correlation (Pearson r = 0.35). Enzymes involved in bioenergetic pathways are indicated in blue and lipid metabolic enzymes in orange. A linear regression line is shown in black. Bottom panels: Global RNA–protein correlations for all detected genes/proteins. The left panel shows the comparison between CSCs and WT cells (Pearson r = 0.54), whereas the right panel shows CSCs treated with HMPSNE compared with untreated CSCs (Pearson r = 0.26). Each point represents a gene/protein pair detected in both datasets. Linear regression lines illustrate the overall correlation trends. These analyses demonstrate that transcriptomic and proteomic changes are moderately correlated in CSCs relative to WT cells, whereas HMPSNE treatment induces broader proteomic remodeling with weaker correspondence between RNA and protein abundance changes.

---

**Extended Data Figure 1.**
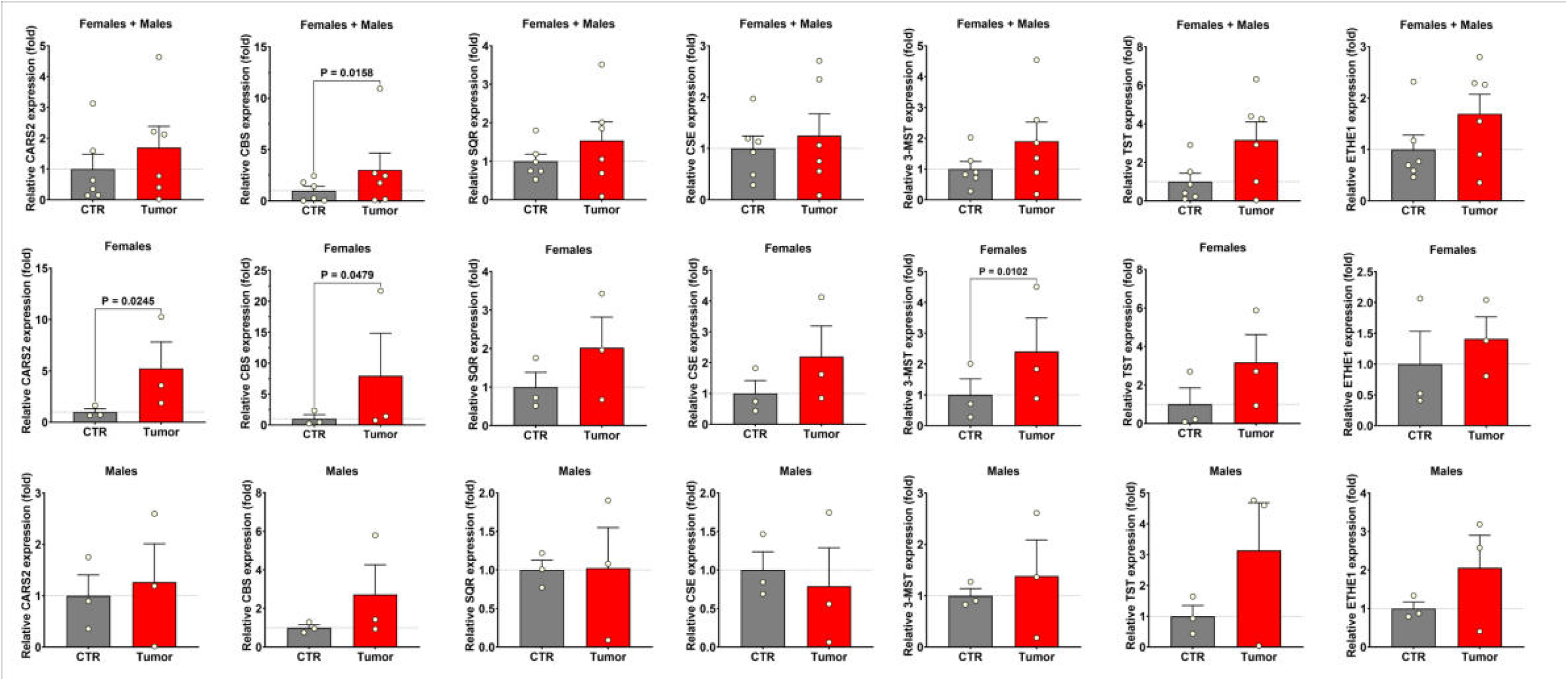
Expression of H_2_S-related enzymes in human colorectal tumor tissue. Relative protein expression of H_2_S-producing and H_2_S-metabolizing enzymes in colorectal tumor samples compared with matched control tissue (CTR). Enzymes analyzed include cysteinyl-tRNA synthetase 2 (CARS2), cystathionine β-synthase (CBS), sulfide:quinone oxidoreductase (SQR), cystathionine γ-lyase (CSE/CTH), 3-mercaptopyruvate sulfurtransferase (3-MST), thiosulfate sulfurtransferase (TST, rhodanese) and ethylmalonic encephalopathy protein 1 (ETHE1), a mitochondrial sulfur dioxygenase. The top row shows combined analysis of all samples, whereas the middle and bottom rows show the same analysis stratified by sex (females and males, respectively). Individual points represent biological samples and bars indicate mean ± s.e.m. Protein expression is shown as fold change relative to matched control tissue. Statistical comparisons between control and tumor tissues are indicated where significant. Total cohort: n = 6 patients (3 females and 3 males).

**Extended Data Figure 2.**
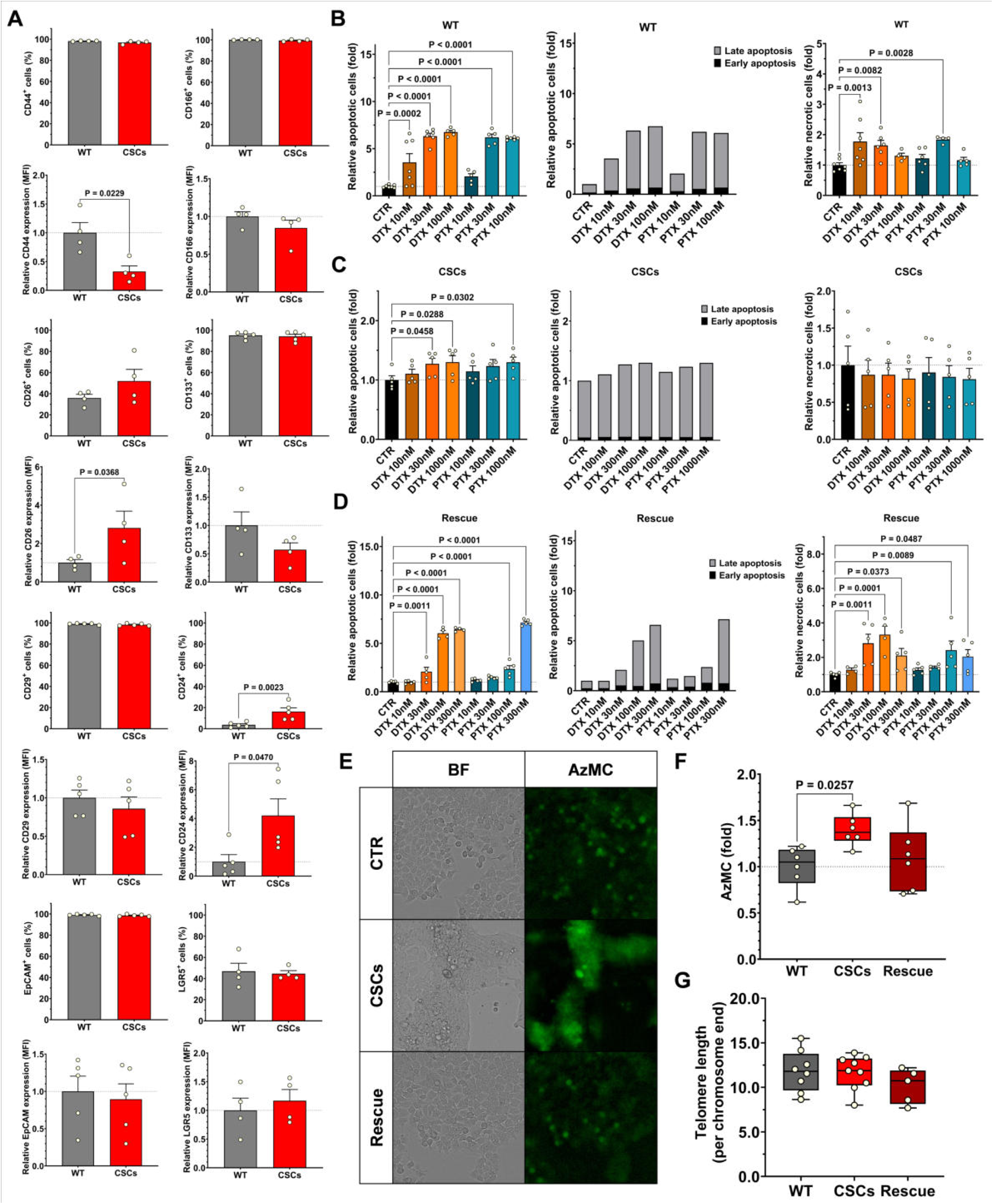
Characterization of colorectal cancer stem cells and their response to chemotherapeutic agents. **(A)** Flow cytometric analysis of stemness-associated surface markers in HCT116 wild-type (WT) cells and cancer stem cells (CSCs). The percentage of positive cells and relative mean fluorescence intensity (MFI) are shown for CD44, CD166, CD26, CD133, CD29, CD24, EpCAM and LGR5. CSCs display altered expression of several stemness-associated markers compared with WT cells. **(B–D)** Quantification of apoptotic and necrotic cell death in WT (B), CSCs (C) and Rescue (D) cells following treatment with the microtubule-targeting chemotherapeutic agents docetaxel (DTX) or paclitaxel (PTX) at the indicated concentrations. Total apoptotic cells are shown together with the distribution of early (black) and late (grey) apoptotic populations, as well as relative necrotic cell levels. CSCs exhibit reduced induction of cell death compared with WT cells even at tenfold higher drug concentrations. **(E)** Representative bright-field (BF) and AzMC fluorescence images of WT, CSCs and Rescue cells showing relative intracellular H_2_S levels. AzMC (7-azido-4-methylcoumarin) fluorescence was used as a probe for H_2_S detection. **(F)** Quantification of AzMC fluorescence intensity (fold change) in WT, CSCs and Rescue cells. **(G)** Telomere length measurements expressed per chromosome end in WT, CSCs and Rescue cells. Data are presented as mean ± s.e.m. from ≥4 independent experiments unless otherwise indicated.

**Extended Data Fig. 3.**
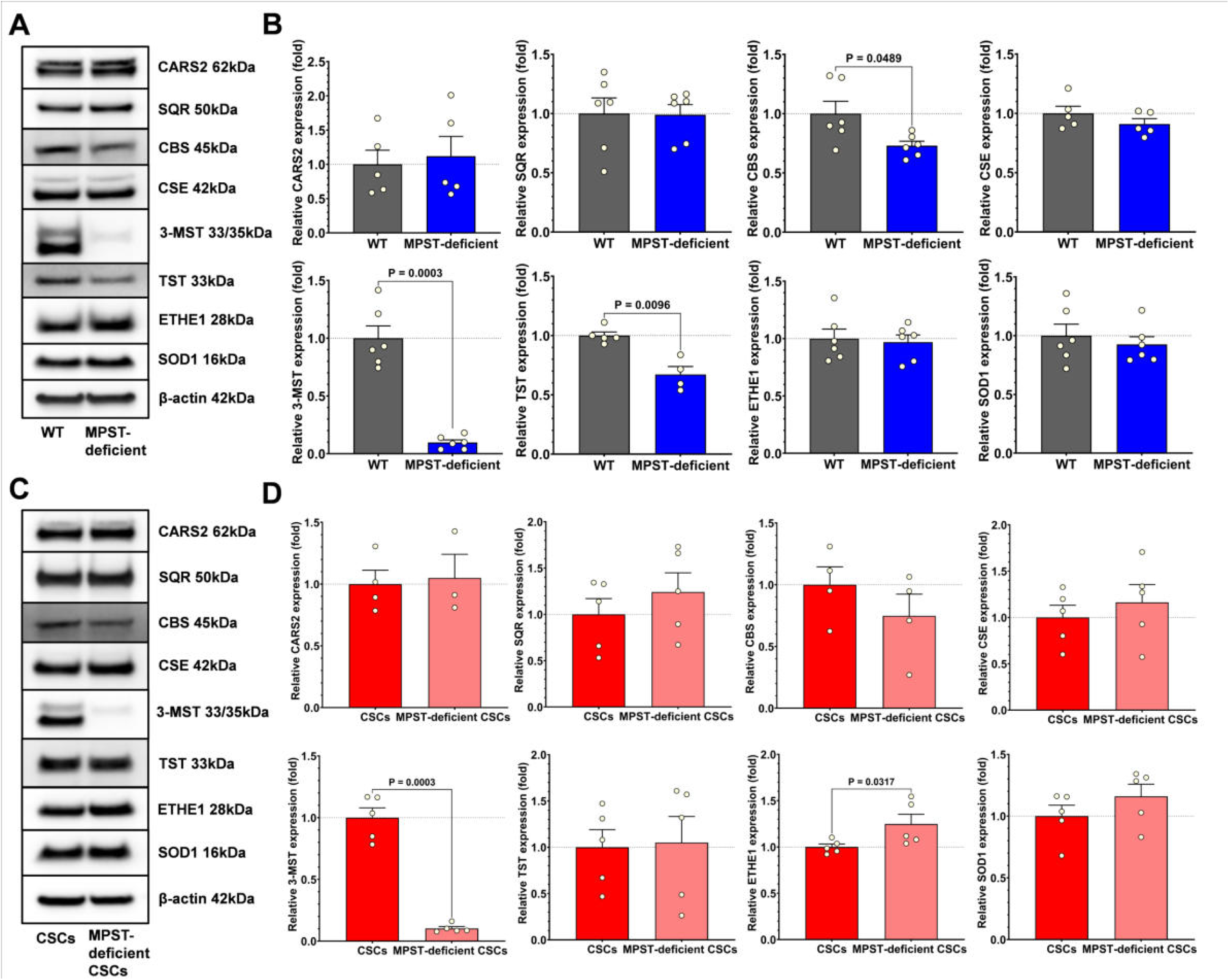
Effects of reduced 3-MST expression on H₂S-related metabolic enzymes. **(A, C)** Representative immunoblots showing protein expression of H_2_S-related enzymes in (A) HCT116 wild-type (WT) and MPST-deficient cells, and (C) cancer stem cells (CSCs) and MPST-deficient CSCs. Proteins analyzed include cysteinyl-tRNA synthetase 2 (CARS2), cystathionine β-synthase (CBS), sulfide:quinone oxidoreductase (SQR), cystathionine γ-lyase (CSE/CTH), 3-mercaptopyruvate sulfurtransferase (3-MST), thiosulfate sulfurtransferase (TST, rhodanese) and ethylmalonic encephalopathy protein 1 (ETHE1). β-actin was used as a loading control. **(B, D)** Densitometric quantification of protein expression in WT and MPST-deficient cells (B), and in CSCs and MPST-deficient CSCs (D). Protein levels were normalized to β-actin and expressed relative to WT or CSC controls, respectively. Reduced 3-MST expression is associated with changes in the expression of selected components of the sulfur metabolism pathway. Data are presented as mean ± s.e.m. from ≥4 independent experiments unless otherwise indicated.

**Extended Data Figure 4.**
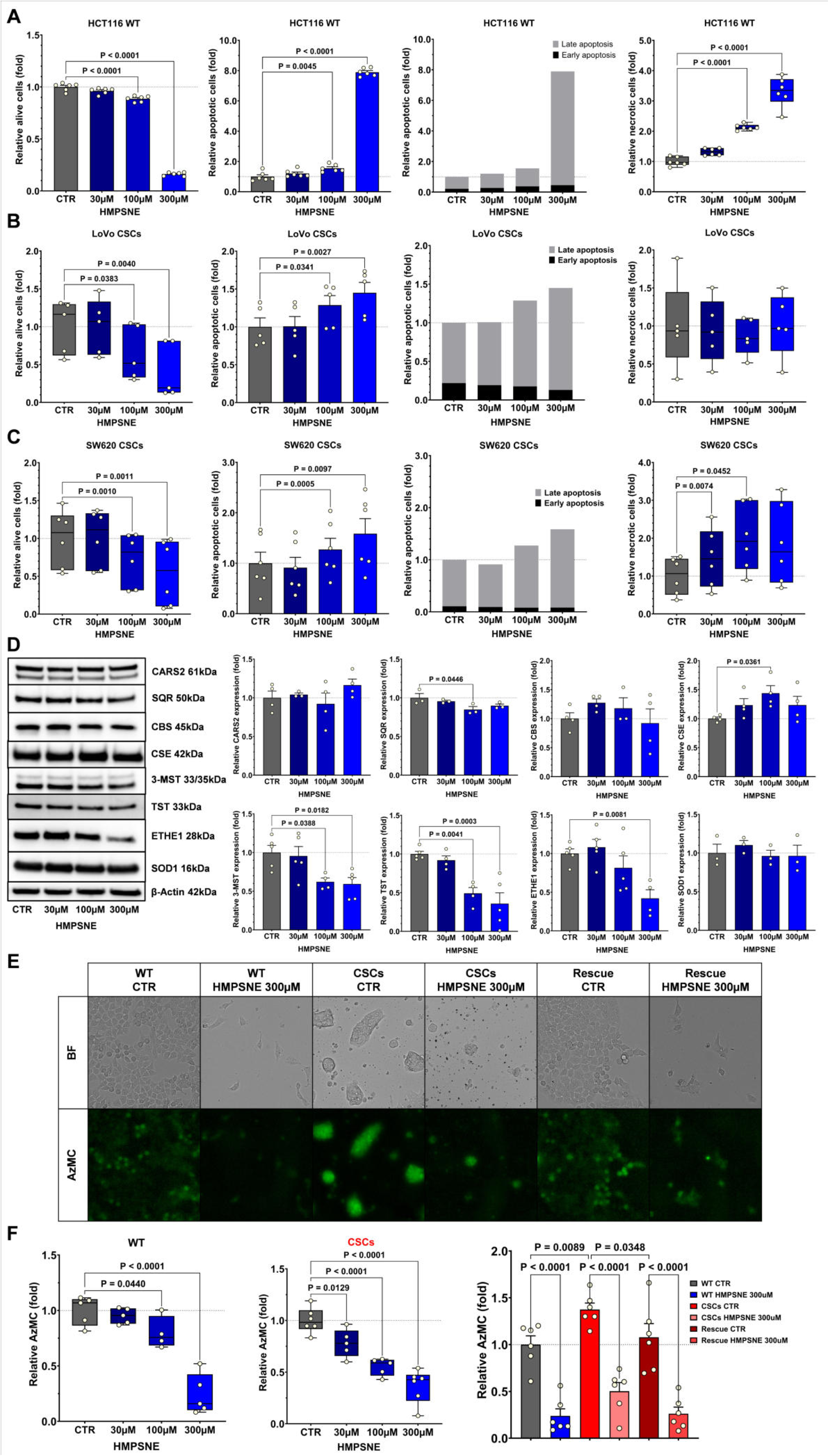
Effects of pharmacological 3-MST inhibition on colorectal cancer cell viability, sulfur metabolism enzymes and H_2_S production. (A–C) Quantification of cell viability, apoptosis and necrosis in HCT116 wild-type (WT) (A), LoVo CSCs (B) and SW620 CSCs (C) treated with increasing concentrations of the 3-MST inhibitor HMPSNE (30–300 μM). Total apoptotic cells are shown together with the distribution of early (black) and late (grey) apoptotic populations, as well as relative necrotic cell levels. HMPSNE treatment reduces cell viability and increases apoptotic and necrotic cell populations in a concentration-dependent manner. Data are presented as mean ± s.e.m. from ≥5 independent experiments. **(D)** Representative immunoblots and densitometric quantification of H_2_S-related metabolic enzymes in HCT116 CSCs treated with increasing concentrations of HMPSNE, including cysteinyl-tRNA synthetase 2 (CARS2), sulfide:quinone oxidoreductase (SQR), cystathionine β-synthase (CBS), cystathionine γ-lyase (CSE), 3-mercaptopyruvate sulfurtransferase (3-MST), thiosulfate sulfurtransferase (TST), ethylmalonic encephalopathy protein 1 (ETHE1) and superoxide dismutase 1 (SOD1). β-actin was used as a loading control. Protein levels were normalized to untreated control (CTR) conditions. Data are presented as mean ± s.e.m. from ≥3 independent experiments. **(E)** Representative bright-field (BF) and AzMC fluorescence images of WT cells, CSCs and Rescue cells treated with HMPSNE (300 μM) showing reduced intracellular H₂S-dependent fluorescence signal following 3-MST inhibition. AzMC fluorescence was used as a probe to detect intracellular H₂S. **(F)** Quantification of AzMC fluorescence in WT cells, CSCs and Rescue cells treated with increasing concentrations of HMPSNE, demonstrating a concentration-dependent reduction in intracellular H_2_S signal following 3-MST inhibition. Data are presented as mean ± s.e.m. from ≥4 independent experiments.

**Extended Data Figure 5.**
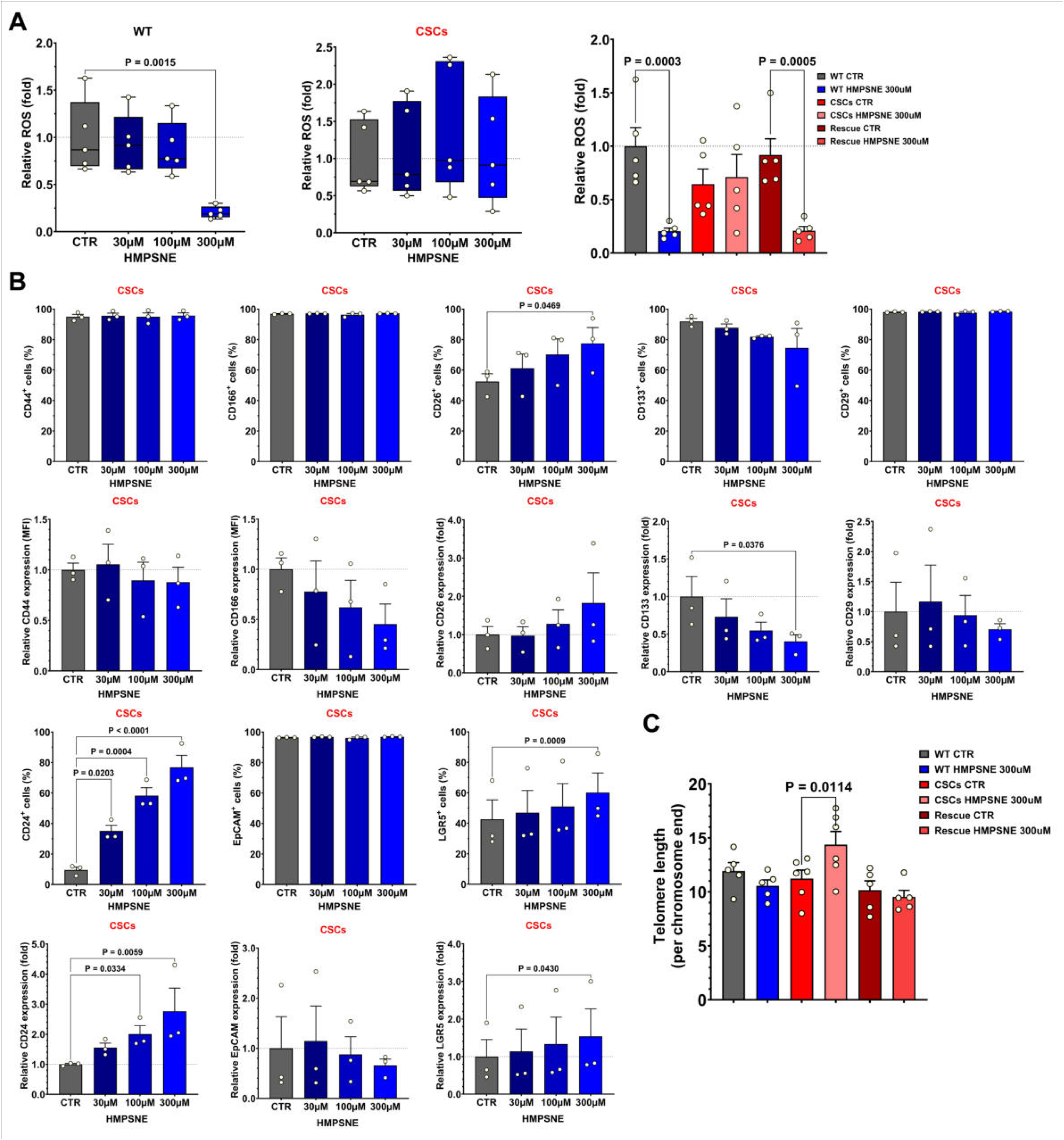
Effects of pharmacological 3-MST inhibition on oxidative stress, stemness markers and telomere length. **(A)** Quantification of intracellular reactive oxygen species (ROS) levels in HCT116 wild-type (WT) cells and cancer stem cells (CSCs) treated with increasing concentrations of the 3-MST inhibitor HMPSNE (30–300 μM). Additional comparison of ROS levels in WT, CSCs and Rescue cells under control (CTR) and HMPSNE-treated (300 μM) conditions is shown. HMPSNE treatment modulates ROS levels in both WT and CSCs populations. **(B)** Flow cytometric analysis of stemness-associated surface markers in HCT116 CSCs treated with increasing concentrations of HMPSNE (30–300 μM). The percentage of positive cells and relative expression levels (mean fluorescence intensity, MFI) are shown for CD44, CD166, CD26, CD133, CD29, CD24, EpCAM and LGR5. HMPSNE treatment alters the expression of several stemness-associated markers. **(C)** Telomere length measurements expressed per chromosome end in WT cells, CSCs and Rescue cells treated with HMPSNE (300 μM). Data are presented as mean ± s.e.m. from 5 independent experiments unless otherwise indicated (B, n = 3).

## Notes

### Competing Interest Statement

The authors have declared no competing interest.

